# miniVec: A miniaturized plasmid backbone supporting antibiotic-free and additive-free fermentation with enhanced yield and functionality

**DOI:** 10.1101/2025.10.02.679775

**Authors:** Silk J. Shi, Yingjun Lin, Jason Z. Ye, Aaron Z. Kwok, Kyrie Z. Wang, Jesse Z. Cai, Mina M. Hu, Arianna Y. Liu, Kyler J. Li, Bernice Y. Guo, Herman H. Xia, Evian P. Huang, Julia X. Chen, Jack F. Hong, Cole K. Zheng, Bruce T. Lahn

## Abstract

Plasmids are a foundational research reagent and a key material in biopharmaceutical manufacturing. Plasmids require a backbone for propagation in E. coli, which typically contains an antibiotic resistance gene alongside a replication origin. As plasmids increasingly enter clinical applications, concerns are raised on the safety risks of antibiotic resistance genes.

Additionally, these protein-coding genes occupy long stretches of DNA that incur significant metabolic burdens on host cells, which negatively impacts plasmid manufacturability and functionality, leading to high production cost and compromised clinical efficacy. Here, we describe miniVec, a novel miniaturized plasmid backbone devoid of protein-coding sequence, and instead expresses a small RNA to provide constant selective pressure capable of sustaining high plasmid copy numbers in plain culture media devoid of antibiotics or other chemical additives. This simplifies large-scale fermentation and greatly increases plasmid yield. Notably, miniVec confers enhanced functionality in a variety of applications such as chemical transfection, electroporation, virus packaging, transposon-or CRISPR-mediated genome integration, and in vivo naked DNA transfection and vaccination, while exhibiting no detectable immunogenicity or toxicity. These advantages establish miniVec as the next-generation plasmid platform for clinical applications, featuring improved safety that aligns with regulatory expectations, enhanced manufacturability leading to much higher yield and dramatic cost reduction, and augmented functionality in diverse applications.

## INTRODUCTION

Plasmids have long been an indispensable tool in gene cloning, manipulation and delivery. Besides being a workhorse of life sciences research, they play an increasingly important role in clinical applications, with either direct human use such as DNA-based vaccines and gene therapy vehicles, or as starting materials for manufacturing various biopharmaceuticals such as recombinant proteins, viral vectors, and mRNA (1–8). Of note is the prominent role that plasmids played in serving as in vitro transcription templates to produce mRNA vaccines to combat the COVID-19 pandemic (9, 10).

Plasmids used in labs are typically extrachromosomal DNA in circular, double-stranded form capable of replicating in E. coli host. Plasmids require a backbone to support their replication and long-term maintenance in host cells, which typically includes an origin of replication (Ori) and an antibiotic resistance gene (ARG). The functional sequence containing the gene of interest (GOI) is placed on the backbone as a hitchhiker.

The ARG allows the use of antibiotics during E. coli fermentation to provide positive selective pressure needed to maintain plasmids in host cells. However, the use of ARGs has raised significant concerns especially in clinical applications. Horizontal gene transfer may create antibiotic-resistant “superbugs”, and residual antibiotics in therapeutic products may provoke hypersensitivity in patients. As such, regulatory authorities such as the FDA and EMA discourage the use of antibiotics in the production of medical-grade plasmids, especially the beta-lactam class of antibiotics (11–15). Additionally, being protein-coding genes, ARGs occupy long stretches of DNA whose replication, transcription and translation place a significant metabolic burden on host bacteria, which negatively impacts plasmid yield and raises production cost (16, 17). Furthermore, when introduced into eukaryotic systems, prokaryotic backbones often form compact, heterochromatin-like structures, leading to silencing of the GOI and unwanted immune response triggered by unmethylated CpG motifs (18–23). Consequently, traditional plasmids as a pharmaceutical material are not only suboptimal in safety and function, but very costly to produce, prohibitively so for some therapeutic applications. As more plasmids move from bench to bedside, the need to address the limitations of traditional plasmid backbones has grown increasingly pressing.

Several plasmid backbone designs attempted to address the above needs (24). The strategy of integrating an E. coli growth inhibitor gene (e.g., ccdB) into the E. coli host genome and placing its antidote gene (e.g., ccdA) on the plasmid backbone provides non-antibiotic selection (25), but still leaves a long protein-coding gene on the backbone that poses safety concerns and incurs a significant metabolic burden. Other antibiotic-free systems, such as Nanoplasmid that employs the toxin gene levansucrase, regulated by a small RNA known as RNA-OUT, to convert sucrose to levan that is toxic to E. coli (26, 27), require additional chemical additives and complex induction protocols during fermentation, which compromises manufacturability by introducing fermentation inconsistencies, purification challenges, and additional quality control (QC) testing for residual additives and their metabolic derivatives. The Minicircle system eliminates antibiotic resistance genes post-fermentation by recombinase and endonuclease (28, 29), but suffers from complex manufacturing protocols and severely reduced plasmid DNA yield. A selection method that is antibiotic-free, additive-free, dispensing with protein-coding genes altogether, while offering easy manufacturing process and high plasmid yield is therefore highly desirable. Additionally, a selection system that minimizes the length of the backbone is greatly preferred as this could enhance plasmid functionality including transgene expression while mitigating episomal silencing and host immune response in eucaryotic cells (30–32). Moreover, during viral vector production, accidental packaging of unwanted sequences from packaging plasmids is a known risk, which further motivates the shift toward miniaturized backbones (33). Indeed, as a matter of good practice, food and drug regulatory authorities such as the FDA and EMA generally emphasize removing non-essential elements in drug design wherever possible to mitigate known or unknown risks they may pose (33, 34).

Here, we report miniVec, the first antibiotic-free, additive-free, non-protein-coding, and miniaturized plasmid backbone system that addresses all the major challenges of traditional backbones more thoroughly than the other approaches. We present data showing that miniVec enables much higher plasmid yield under simpler manufacturing process in both lab and industrial settings. Furthermore, the system significantly enhances plasmid functionality in a variety of applications such as chemical transfection, electroporation, virus packaging, transposon-or CRISPR-mediated genome integration, and in vivo naked DNA transfection and vaccination, while exhibiting no detectable immunogenicity and showing an excellent safety profile in vivo. For gene therapy in particular, the outstanding performance of miniVec in plasmid production and downstream manufacturing of viral and nonviral vectors creates tremendous compound cost savings for this budget-unfriendly sector. Our results thus establish miniVec as the next-generation plasmid platform for clinical applications especially in the manufacturing of genetic medicines.

## RESULTS

### Design principle of miniVec

A key innovation of the miniVec system lies in its ability to support antibiotic-free and additive-free maintenance of plasmids in a specially engineered E. coli host, the miniHost (Figure 1). As such, only regular growth media devoid of any antibiotics or other chemical additives is sufficient for the proliferation of miniVec plasmids, which is particularly advantageous in large-scale manufacturing of clinical-grade plasmid DNA by good manufacturing process (GMP).

**Figure 1.**
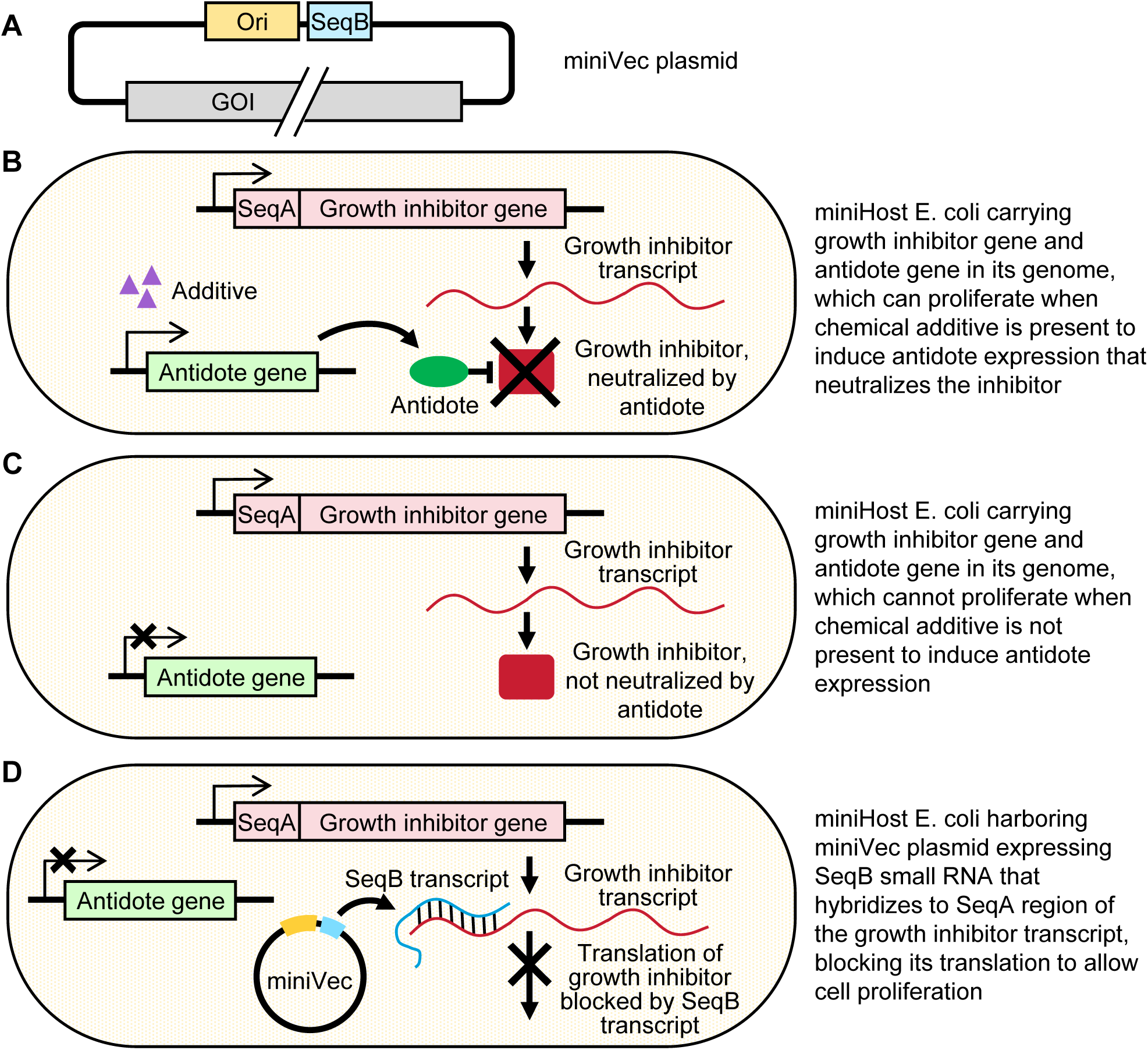
Schematic of antibiotic-free and additive-free selection of the miniVec plasmid in the specially engineered E. coli host miniHost. **(A)** Structure of the miniVec backbone, which contains the small RNA gene SeqB alongside an Ori. **(B)** The genome of miniHost carries one or more copies of a growth inhibitor gene with a sequence known as SeqA embedded in its 5’ UTR. The protein product of this gene blocks host cell proliferation. The miniHost genome also carries an antidote gene whose protein product can neutralize the growth inhibitor to allow cells to proliferate. Expression of the antidote is inducible by a chemical additive. For miniHost cells not yet harboring any miniVec plasmid, this additive is added in the media to induce expression of the antidote to neutralize the growth inhibitor in order for cells to grow. **(C)** Upon transformation of miniHost by the miniVec plasmid, the additive is removed such that the antidote is no longer expressed. For cells that fail to take in the miniVec plasmid, proliferation is blocked by the growth inhibitor because the antidote is no longer present. **(D)** The SeqB small RNA gene on the miniVec plasmid backbone produces a small noncoding RNA complementarity to the SeqA region of the growth inhibitor transcript, such that the SeqB RNA can hybridize to SeqA. This, plus additional secondary structure on the SeqB RNA, inhibits the translation of the growth inhibitor to allow cell proliferation. Upon transformation of miniHost by the miniVec plasmid and removal of the additive, cells successfully transformed by the miniVec plasmid can proliferate due to neutralization of the growth inhibitor gene in the miniHost genome by the SeqB small RNA expressed from the miniVec backbone.

The miniVec backbone carries a small RNA gene known as SeqB alongside an Ori (Figure 1A). The genome of miniHost carries one or more copies of a growth inhibitor gene with a sequence known as SeqA embedded in its 5’ UTR. This growth inhibitor gene, when properly expressed, blocks proliferation of the E. coli host. The genome of miniHost also carries an antidote gene whose protein product can neutralize the growth inhibitor. Expression of the antidote gene is inducible by a chemical additive. For the miniHost E. coli not yet harboring any plasmid, this additive is needed in the culture media to induce expression of the antidote gene that in turn neutralizes the growth inhibitor to allow cell proliferation (Figure 1B). Importantly, upon transformation of miniHost by a miniVec plasmid, the additive is removed. Cells that fail to take in the miniVec plasmid cannot proliferate due to the presence of the growth inhibitor and the absence of antidote expression without the additive (Figure 1C). In cells transformed by the miniVec plasmid, the SeqB gene on the miniVec backbone produces a small noncoding RNA complementary to the SeqA region of the growth inhibitor transcript, such that the SeqB RNA can hybridize to SeqA, suppressing translation of the growth inhibitor, which in turn allows cells to proliferate (Figure 1D). The SeqB RNA can also contain sequences that form secondary structures to further inhibit the translation of the inhibitor gene by blocking the loading and/or processing of ribosomes on the inhibitor gene transcript. Consequently, the miniVec backbone only contains two short functional components – the SeqB small RNA gene alongside an Ori that supports plasmid replication in E. coli. The backbone is completely devoid of any protein-coding sequence and, depending on the Ori selected, can be as short as ∼500bp, which incurs minimal metabolic burden on the host.

By the above design, the miniVec plasmid is under constant selection to be present in miniHost cells grown in regular media devoid of any antibiotics or other additives. Furthermore, selective pressure favors high copy numbers of the plasmid in cells to better counter the deleterious effect of the growth inhibitor on cell proliferation, which contributes to exceptionally high plasmid DNA yield as described below. In principle, selective pressure for high plasmid copy number can be raised further by increasing the copy number of the integrated growth inhibitor gene and tweaking SeqA and SeqB sequences to optimize the translational suppression of the growth inhibitor.

We tested a number of growth inhibitor and antidote gene pairs as well as a variety of SeqA and SeqB pairs that supported the intended plasmid maintenance in host cells. In this study, we focused on miniHost strains that carried one or more genomic copies of the ccdB gene that inhibits E. coli growth, and the ccdA antidote gene derived from the F plasmid (35, 36), with ccdA designed to be inducible by arabinose, and SeqA and SeqB sequences derived from the small-RNA-based negative autoregulation system of the R1 plasmid (37), along with shortened pUC Ori or R6K Ori.

We have successfully built a large number of plasmids on the miniVec backbone that varied in size, GOI (including both protein coding genes and small RNA genes such as gRNA and shRNA genes), and GC content. The cloning efficiency was comparable to traditional antibiotic resistance backbones. To date, the largest construct that we have successfully created using the miniVec backbone was an adenoviral packaging plasmid with a total size of 34,865 bp, though the upper limit of miniVec remains open as we have not yet attempted big plasmids.

### Long-term stability of miniVec plasmid

To test the long-term stability of miniVec plasmids in miniHost, duplicates of three different miniVec plasmids including AAV EGFP transfer plasmid, and plasmids expressing VSV-G or lentiviral GagPol were passaged for 15 days consecutively with approximately 250 total doublings. On day 11 and day 15, 96 colonies were picked from each duplicate of each plasmid, and tested for the presence of the plasmid. All 96 clonal colonies from each sample showed the presence of the plasmid at both timepoints, indicating a 100% plasmid retention rate. Full length sequencing was performed on 10 out of the 96 colonies from day 15. All resulting sequences aligned perfectly with the reference sequence from day 0, indicating sequence stability. The miniVec system thus shows excellent long-term stability.

### Substantially increased DNA yield of miniVec plasmid

Plasmid DNA yield of the miniVec backbone was compared to the traditional antibiotic resistance backbone in lab-scale culture using Erlenmeyer shake flasks. The miniVec version and the traditional version were each tested with 5 inserts, including AAV EGFP transfer plasmids, lentivirus EGFP transfer plasmids, EGFP-expressing transposon plasmids including PiggyBac and Sleeping Beauty, as well as lentiviral packaging Rev plasmids, totaling 10 plasmids. After culturing in shake flasks overnight, plasmid DNA was isolated from bacterial culture by standard miniprep. Comparing the volumetric yield under the same condition, a substantial increase for the various plasmids was observed by simply switching the backbone from the traditional version to miniVec (Figure 2A). The highest yield in shake flask was 10.97 mg per liter of bacterial culture on average for the PiggyBac EGFP transposon plasmid on miniVec backbone compared to 5.78 mg per liter of its traditional backbone counterpart. The higher yield of the miniVec backbone is even more pronounced when measuring plasmid yield by copy number rather than mass (Figure 2B). The miniVec PiggyBac EGFP transposon plasmid achieved 5.28 nmol per liter on average compared to the 1.56 nmol per liter of its traditional counterpart.

**Figure 2.**
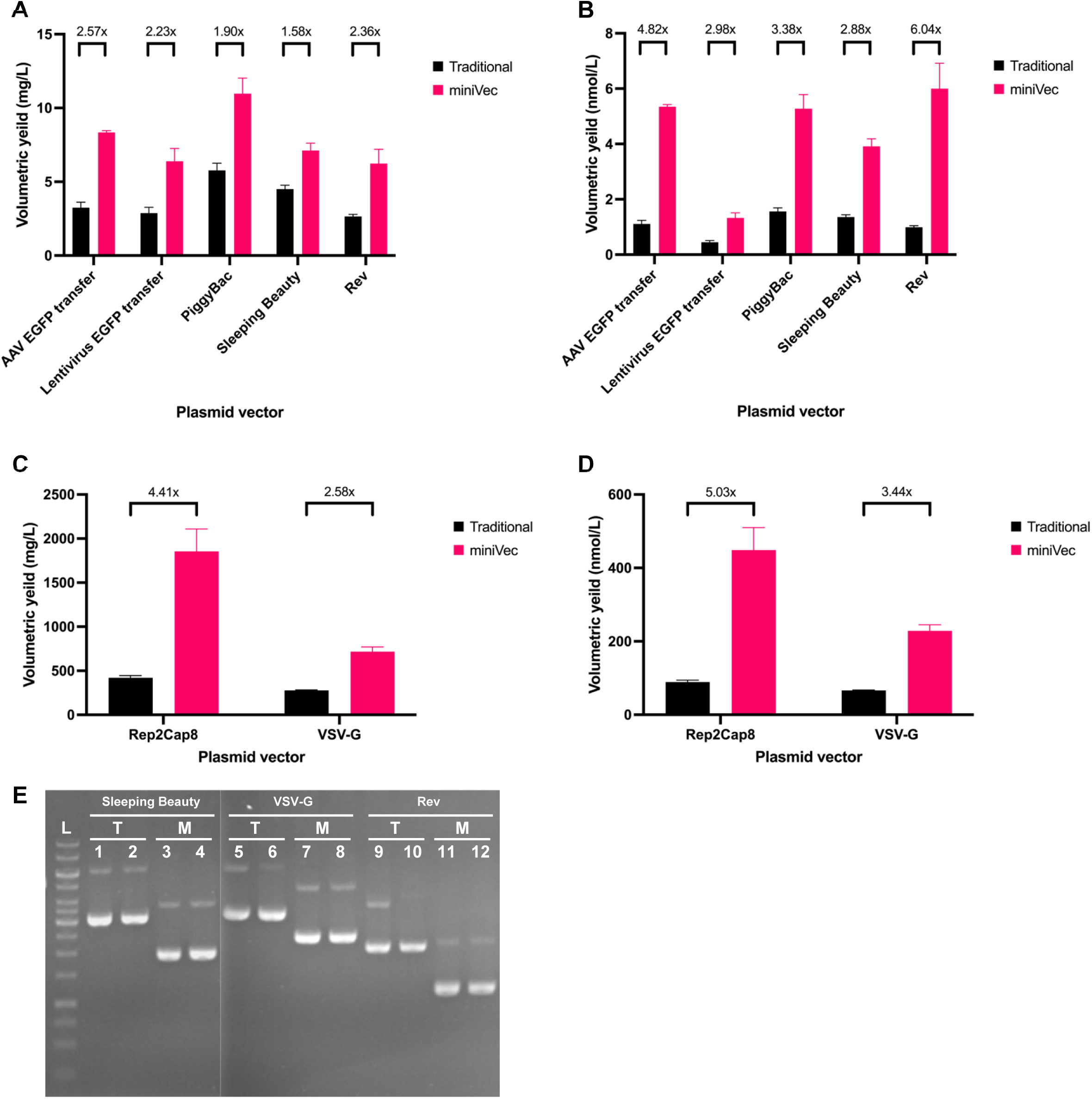
Comparison of plasmid DNA yield between miniVec and traditional backbones. **(A, B)** E. coli carrying each plasmid was grown in lab-scale Erlenmeyer shake flasks, and plasmid yield per unit volume of E. coli culture is given in mass **(A)** or molar quantity **(B)**. E. coli was grown in a 2.7-liter fermenter used in industrial-scale manufacturing, and yield per unit volume of E. coli culture is given in mass **(C)** or molar quantity **(D)**. **(E)** Homogeneity of plasmid configuration was tested by gel electrophoresis. Even number lanes were samples treated with T5 exonuclease that digests away all but double-stranded covalently closed circular DNA (cccDNA), while odd number lanes were untreated controls. T: traditional backbone; M: miniVec backbone.

The miniVec and traditional plasmids were also compared for yield in industrial fermentation. The scale was 2.7 L using parameters from our platform fermentation technology. Two pairs of plasmids were tested, including AAV packaging Rep2Cap8 plasmids and lentiviral packaging VSV-G plasmids. For each pair, consisting of the same insert on either miniVec or traditional backbone, the fermentation process remained the same for both plasmids except harvest time, which was optimized individually for each plasmid as is typical in industrial fermentation to achieve the best DNA yield. Plasmid DNA was then isolated by standard miniprep to assess volumetric yield. In general, the miniVec backbone brings massive improvement to the volumetric yield (Figure 2C, D). For one comparison, the miniVec plasmid exhibited a yield of 1855.37 mg or 448.45 nmol per liter of harvest, severalfold higher than the 420.80 mg or 89.16 nmol per liter for its traditional counterpart.

The miniVec and traditional plasmids were then tested for the homogeneity of DNA configuration by gel electrophoresis. Three pairs of plasmids were examined, including EGFP-expressing Sleeping Beauty transposon plasmids, and VSV-G and Rev plasmids used in lentivirus packaging. Each pair of plasmids had the same insert on either miniVec or traditional backbone. After culturing in shake flasks, plasmid DNAs were extracted by standard midiprep and treated with T5 exonuclease followed by gel electrophoresis (Figure 2E). T5 exonuclease digests all the single-stranded DNA as well as DNA with an open end. Hence, only plasmids in the form of double-stranded covalently closed circular DNA (cccDNA) can resist T5 digestion. We note that the term “supercoiled DNA” is often used to describe plasmids. We consider “cccDNA” to be the more accurate term as it depicts a definitive state of plasmid configuration that is physically intact – and presumably also functionally intact – given the lack of nicks or broken ends anywhere in the sequence. In contrast, supercoiled DNA is an ambiguous term, both subjectively (i.e., it is unclear just how many positive or negative added coils qualify as supercoil) and objectively (i.e., different buffer conditions can lead to different degrees of coiling, such as high salt increasing coiling and DNA intercalators decreasing coiling). We therefore argue that “cccDNA” should be used as a precise term to describe intact plasmids of any degree of coiling, while “supercoiled DNA” should be used only as an approximate term to refer to the subspecies of cccDNA that contains a large number of positive or negative added coils under regular buffer conditions. In Figure 2E, even number lanes were samples treated with T5 exonuclease while odd number lanes were untreated controls. The assay showed that all miniVec plasmids existed as pure cccDNA given that there is no change in banding patterns from T5 digestion.

### Enhanced chemical transfection and electroporation efficiency of miniVec plasmid

Transient chemical transfection efficiency of miniVec plasmids was compared to the traditional backbone. HEK293T cells were chemically transfected in an equal molar manner with plasmids containing an EGFP overexpression cassette, placed on either the miniVec backbone or the traditional counterpart. Fluorescence imaging revealed brighter EGFP expression of the miniVec plasmids at 48h and 96h post-transfection (Figure 3A). Quantitation by flow cytometry showed that while the rate of fluorescence-positive cells was similar between miniVec and traditional plasmids, mean fluorescence intensity (MFI) was more than 30% higher for miniVec (Figure 3B). The amount of plasmid DNA entering the nuclei of cells may have even greater difference between miniVec and the traditional plasmid, given that the expression of EGFP can reach plateau in cells carrying high copy numbers of the plasmid.

**Figure 3.**
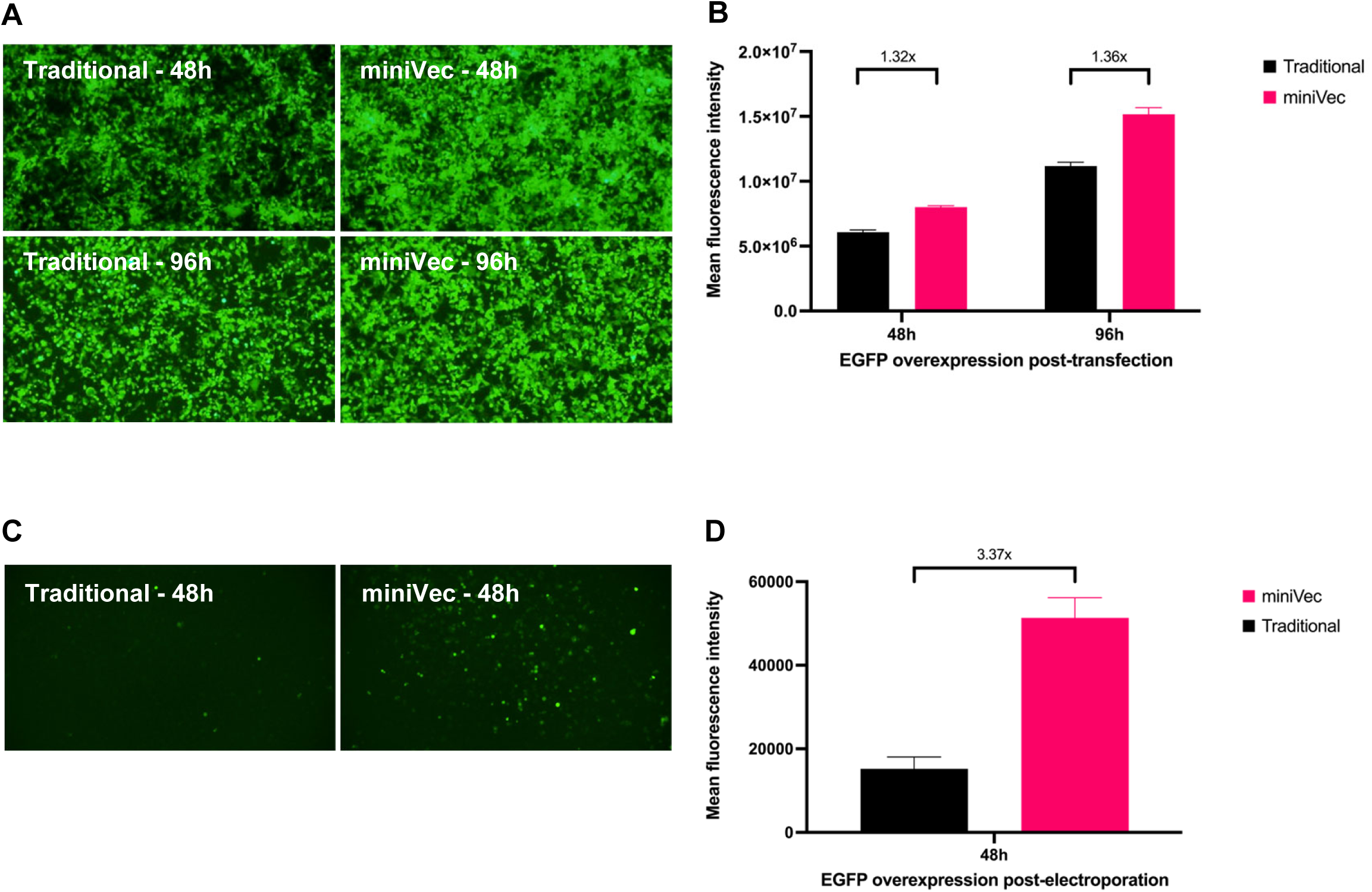
Comparison of chemical transfection and electroporation efficiency between miniVec and traditional plasmids. HEK293T cells were chemically transfected with EGFP overexpression plasmids, either miniVec or traditional, in an equal molar manner, and fluorescence images **(A)** and flow cytometry measurements of mean fluorescence intensity **(B)** were taken at 48h and 96h. Jurkat cells were electroporated with either miniVec or traditional plasmids that carried a CAR-T2A-EGFP bicistronic expression cassette encoding an upstream chimeric antigen receptor (CAR) and a downstream EGFP reporter separated by a T2A, in an equal molar manner, and fluorescence images **(C)** and flow cytometry measurements of mean fluorescence intensity **(D)** were taken at 48h and 96h.

Chemical transfection works poorly on some cell types, in which case electroporation is often employed as an alternative. One potential real-life application is CAR-T where plasmid DNA is delivered into T cells by electroporation to genetically modify the cells. To mimic this application, Jurkat cells, a T lymphocyte cell line, were electroporated with either miniVec or traditional plasmids that carried a CAR-T2A-EGFP bicistronic expression cassette encoding an upstream chimeric antigen receptor (CAR) and a downstream EGFP reporter separated by a T2A, in an equal molar manner. Fluorescence imaging of the electroporated cells after 48h of culture showed dramatically enhanced electroporation efficiency of the miniVec plasmid compared to the traditional one (Figure 3C), with the mean fluorescence intensity as assessed by flow cytometry being several times higher for miniVec (Figure 3D).

### Markedly Increased lentivirus packaging yield with miniVec plasmid

The third-generation packaging system for recombinant lentivirus involves the transient transfection of four plasmids into HEK293T cells, including the lentiviral transfer plasmid carrying the GOI, and the GagPol, Rev, and VSV-G helper plasmids (38). We compared the packaging efficiency of the plasmids that were either all on the miniVec backbone or all on the traditional backbone. The transfer plasmid carried an mCherry reporter gene. The four packaging plasmids, miniVec or traditional, were transfected into HEK293T cells to produce recombinant lentivirus, which was harvested as supernatant at 48h post transfection. Equal volume of each virus was then used to transduce HEK293T cells. Fluorescence imaging indicated higher intensity from the miniVec group (Figure 4A). Measurements by qPCR performed on genomic DNA of transduced cells showed that the titer of the virus packaged with miniVec plasmids was several times higher than traditional plasmids (Figure 4B).

**Figure 4.**
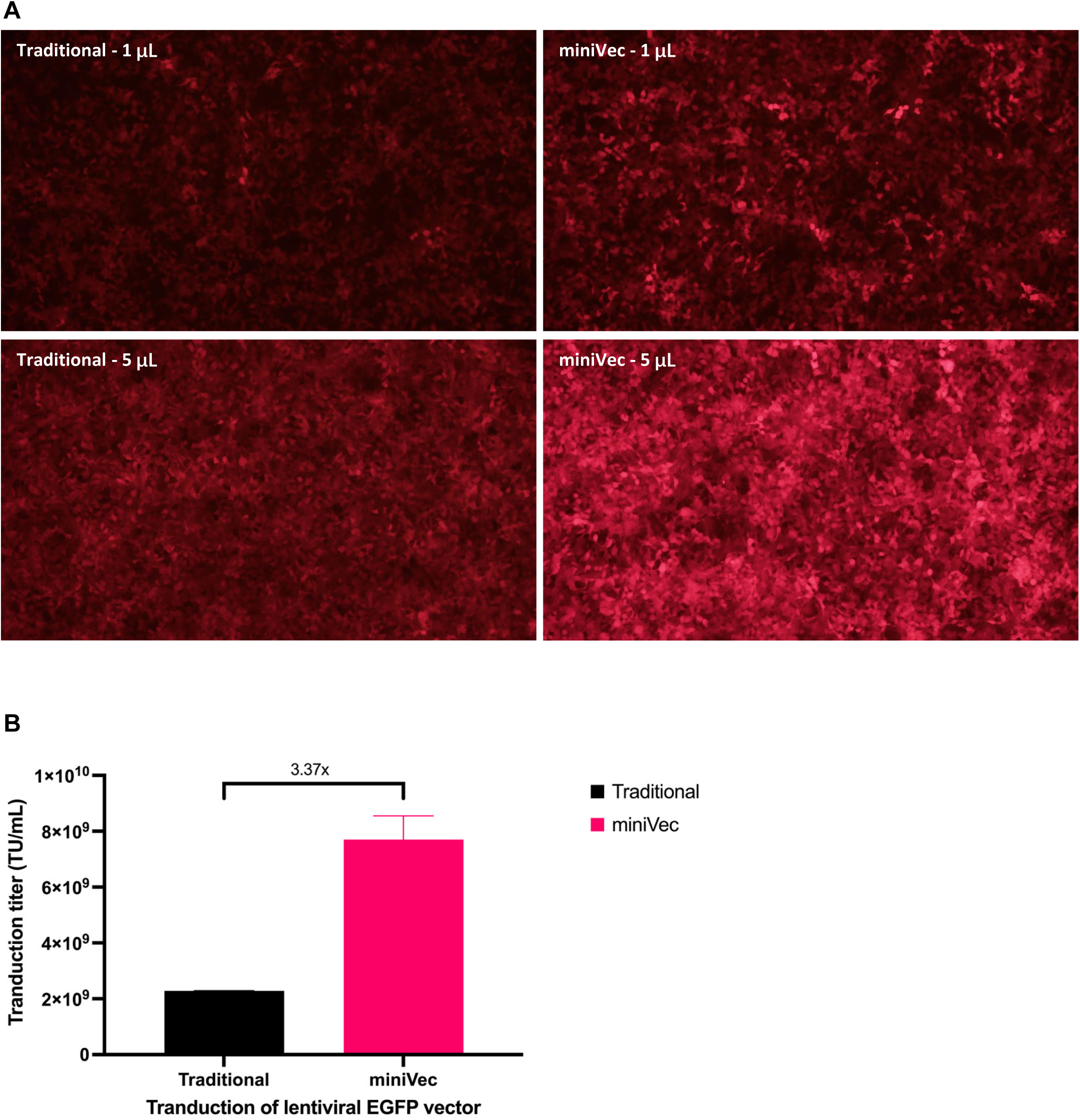
Comparison of functional titer between lentivirus packaged with miniVec plasmids versus traditional plasmids. **(A)** Fluorescence images of HEK293T cells transduced with either 1 μL or 5 μL of lentivirus supernatant packaged from either miniVec or traditional plasmids. **(B)** Titer measurement by qPCR performed on genomic DNA of cells from 0.1, 0.25, and 2.5 μL transduction samples.

### Enhanced efficiency of transposon-mediated genome integration using miniVec plasmid

Transposon systems including piggyBac (PB) and Sleeping Beauty (SB) were examined for how the miniVec backbone affected their integration efficiency. Two separate plasmids were employed in each transposon system, one being the transfer plasmid carrying the transposon that expresses an EGFP reporter, the other being the helper plasmid expressing the PB or SB transposase. For either system, both transfer and helper plasmids were placed on either the miniVec backbone or the traditional backbone. HEK293T cells were transiently co-transfected with the transfer and helper plasmids, either miniVec or traditional, in an equal molar manner between the two backbones. As a control, the transfer plasmid on traditional backbone was also transfected alone without the helper plasmid. Two days after transfection, cells were strongly fluorescent as would be expected for a typical transient transfection that introduced large amounts of episomal plasmid DNA into host cells. Cells were passaged extensive until the control sample had negligible fluorescence left, at which point it was assumed that all the episomal transfer plasmid DNA had been diluted out. Cytometric analysis was then performed to measure EGFP expression that presumably came from the integration of the transfer plasmid into the host genome by transposition. For both PB and SB, samples transfected with miniVec plasmids showed higher levels of fluorescence (Figure 5), indicating that the miniVec backbone increased the efficiency of transposon-mediated genome integration relative to the traditional backbone under equal molar conditions.

**Figure 5.**
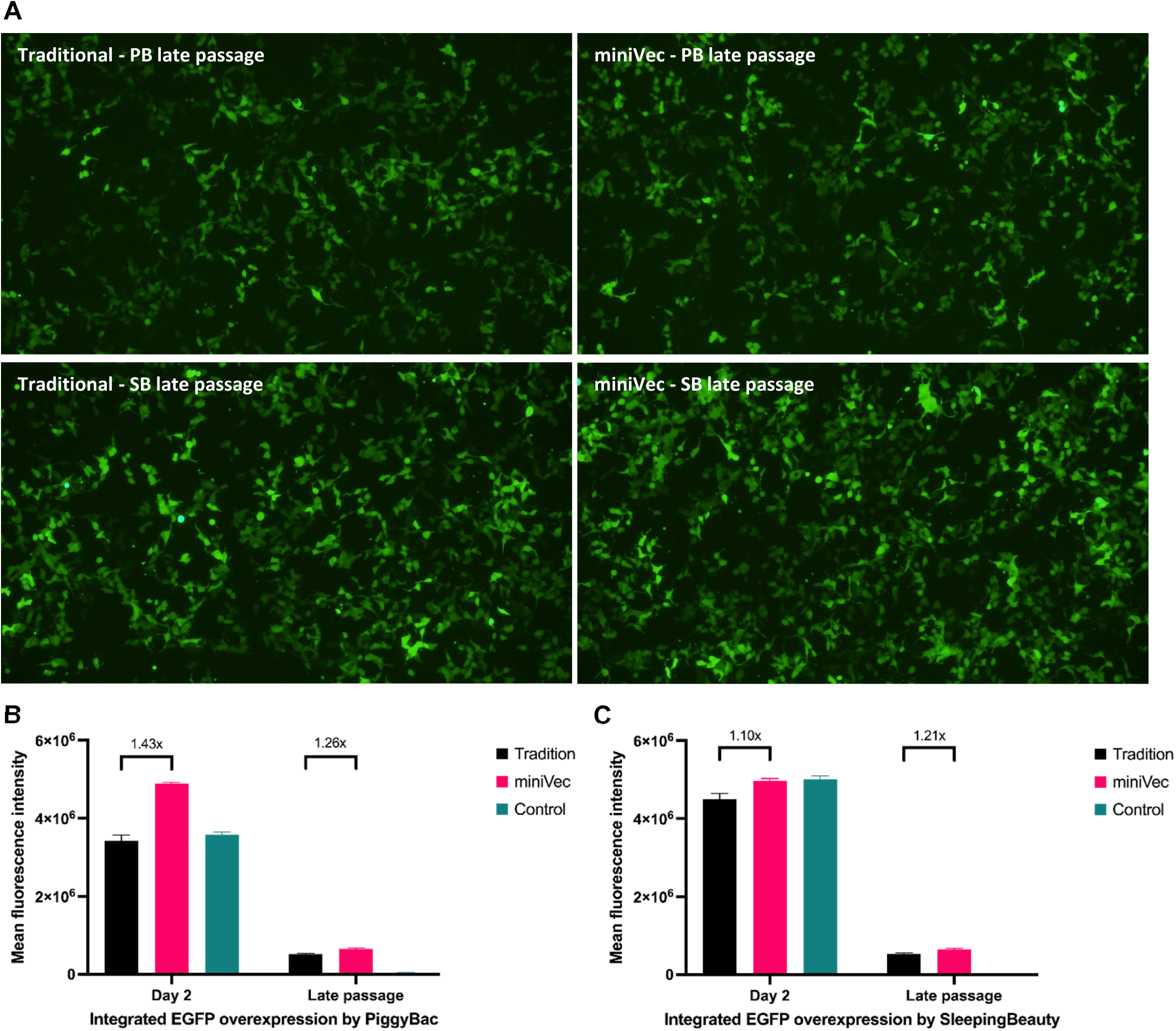
Comparison of transposition efficiency of piggyBac (PB) or Sleeping Beauty (SB) system between miniVec and traditional backbone. Fluorescence images **(A)** were taken for both PB and SB systems on either miniVec or traditional backbone at late passage when the control sample had negligible fluorescence left. Mean fluorescence intensity was measured by flow cytometry at day 2 and late passage for both PB **(B)** and SB **(C)** systems.

In actual transposition applications, the efficiency is typically limited by the upper limit of the DNA mass that can be used in transient transfections. With smaller plasmid size and hence lower molecular weight, the miniVec system could further enhance transposition efficiency by allowing more DNA molar quantity to be used at the mass limit. Indeed, when transfection was done under equal mass condition between miniVec and traditional plasmids, a larger increase in integration-positive cells and fluorescence intensity was observed (data not shown).

While the great majority of integration events should be due to transposition that inserts the transposon into the genome without the plasmid backbone, transposase-independent random integration may still occur at a low frequency that brings the backbone into the host genome. In this case, the miniVec backbone offers the additional advantage of carrying a limited amount of prokaryotic sequence devoid of any protein-coding gene.

### Enhanced efficiency of CRISPR-mediated genome integration using miniVec plasmid

To examine the efficiency of CRISPR-mediated genome integration, we designed hCas9 overexpression plasmid and a donor plasmid expressing EGFP, either miniVec or traditional, for homology-independent targeted insertion (HITI). The target of integration was the AAVS1 locus in HEK293T cells. Thus, the gRNA and cutting sites flanking the knockin cassette on the donor plasmid were designed based on the AAVS1 target. HEK293T cells were transfected with the hCas9 and donor plasmids, either miniVec or traditional, under equal molar conditions. As a control, the donor plasmid was transfected alone without the hCas9 plasmid. Cells were passaged extensively until negligible fluorescence was left in the control sample, which indicated that all the episomal plasmid DNA should have been diluted out. EGFP expression in the HITI samples, which presumably came from the integration of the donor plasmid into the host genome, was measured by flow cytometry. It showed that the miniVec plasmids produced 78% more fluorescence (Figure 6), indicating that the efficiency of HITI-mediated genome integration was much higher for the miniVec backbone relative to the traditional backbone under equal molar conditions.

**Figure 6.**
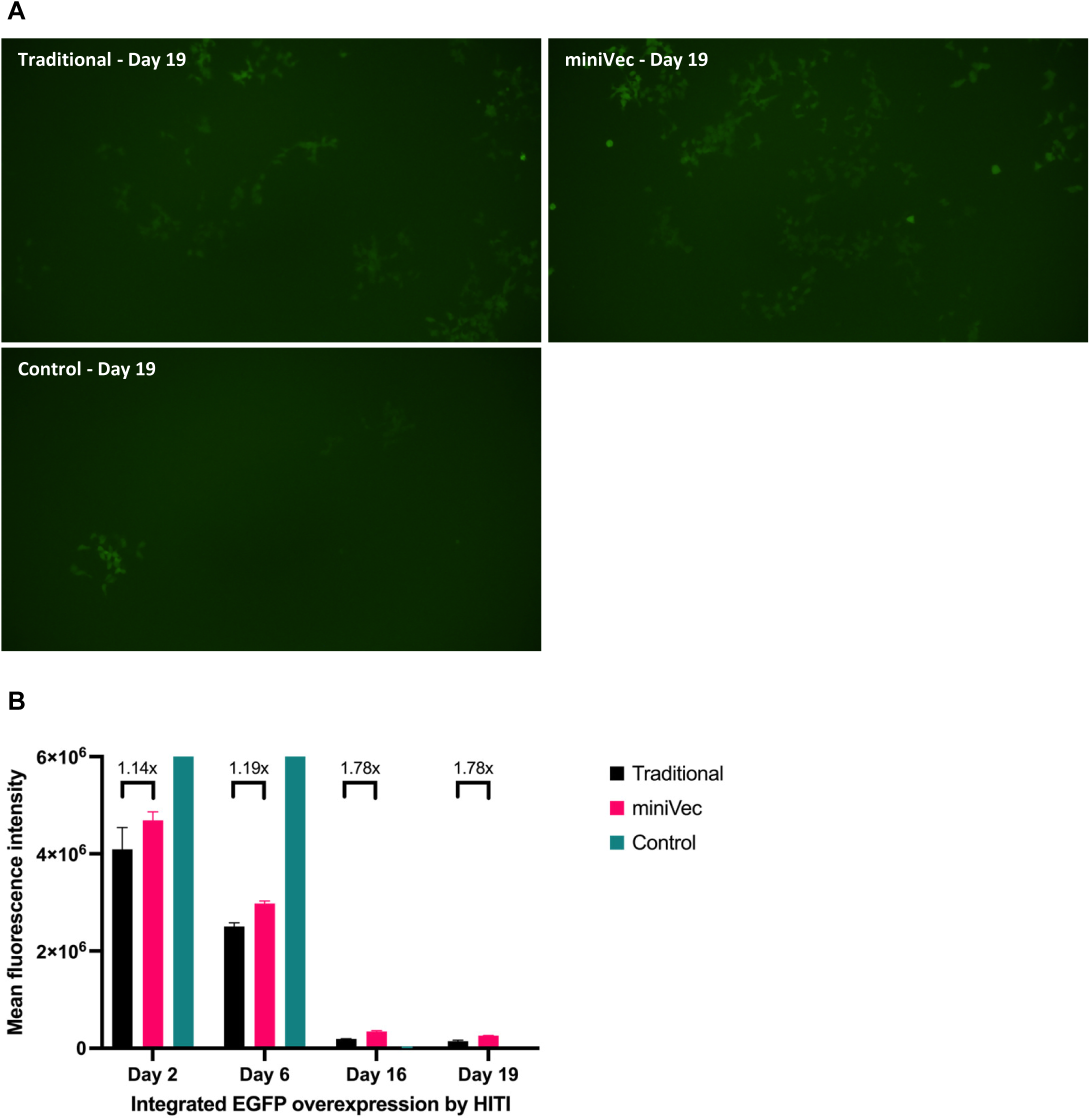
Comparison of HITI efficiency between miniVec backbone and traditional backbone. Fluorescence images **(A)** of the EGFP integrated cells were taken at day 19. Mean fluorescence intensity **(B)** was measured by flow cytometry.

### Enhanced in vivo transfection efficiency using naked miniVec plasmid

The miniVec system was then investigated for its in vivo transfection efficiency. A luciferase vector, either miniVec or traditional in an equal molar manner, was administered as naked DNA by intravenous (IV) or intramuscular (IM) injection into mice. Bioluminescence was measured at several time points post injection. For IV injection, signal came primarily from the liver and subsided after day 7, with the overall luciferase expression several times higher from the miniVec vector relative to the traditional vector on multiple time points (Figure 7A, B). For IM injection, bioluminescence came from the quadricep injection site and lasted much longer before subsiding after 21 days, with severalfold higher luciferase expression from the miniVec vector than the traditional vector on multiple days just like IV injection (Figure 7C, D).

**Figure 7.**
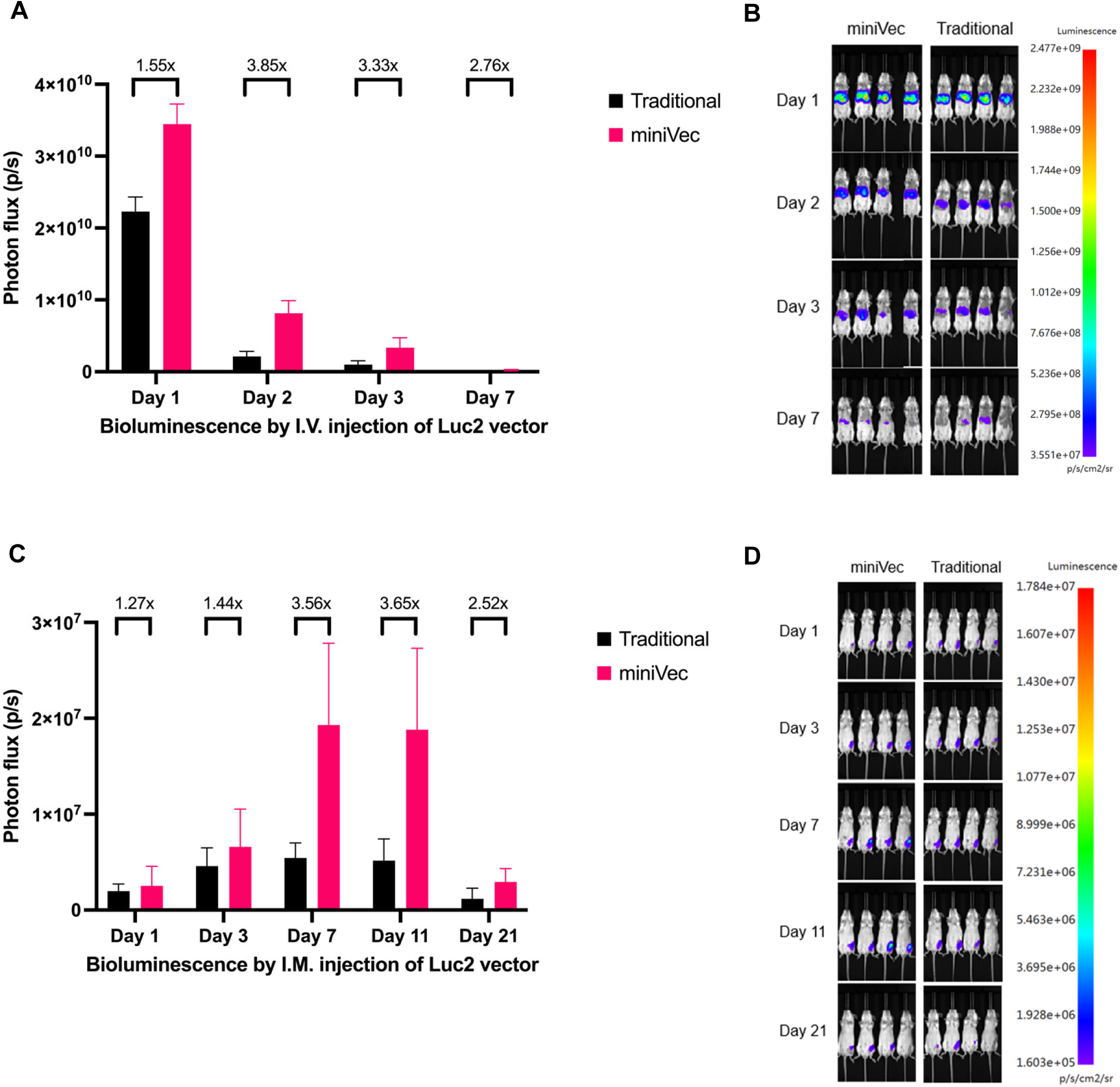
Comparison of in vivo transfection efficiency between miniVec and traditional plasmids. Intravenous (IV) injection through the tail vein **(A, B)**, or intramuscular (IM) injection in the quadricep **(C, D)**, of a plasmid expressing the luciferase report Luc2, on either the miniVec backbone or the traditional backbone in an equal molar manner, were performed. Bioluminescence from luciferase expression was measured on live animals as photon flux (p/s).

Given that the mass of injected DNA is a limiting factor for in vivo transfection, the smaller size of miniVec plasmids should offer even greater advantage over the traditional plasmid when applied with an equal DNA mass.

### Enhanced naked DNA vaccination using miniVec plasmid

Vaccination by naked plasmid DNA is a promising technology with growing applications in infectious disease prevention and the immunotherapy of a range of noninfectious conditions such as cancer, allergy and autoimmune disease (39, 40). It is also an important vaccine modality in veterinary medicine due to its low cost (41–45). With the observation of enhanced in vivo transfection efficiency by naked miniVec plasmid DNA, we wished to study if the efficacy of naked DNA vaccination could be enhanced by the miniVec system. We immunized mice with a plasmid expressing the COVID-19 spike protein, on either miniVec of traditional backbone, with three doses of intramuscular injections. Empty plasmids and PBS were used as negative controls. ELISA was used to detect anti-spike IgG antibody in animals’ sera, which showed higher antibody titers provoked by the miniVec plasmid than the traditional one after each injection, while the controls produced no detectable anti-spike activity (Figure 8A).

**Figure 8.**
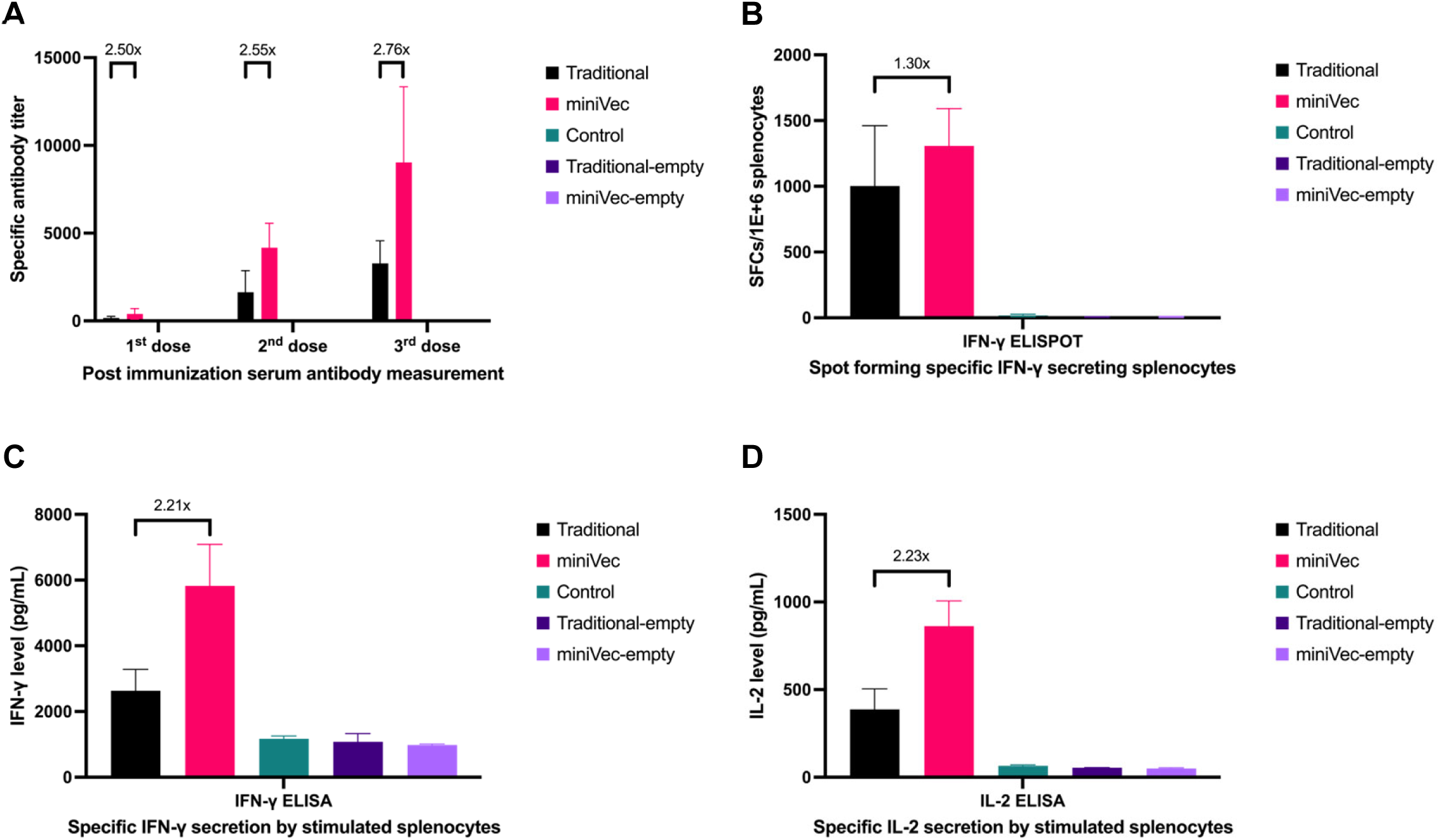
Comparison of immune response to naked DNA vaccination between miniVec and traditional plasmids expressing the COVID-19 spike protein. **(A)** Measurement of anti-spike antibody provoked by naked plasmid DNA vaccination. Retro-orbital blood collection was performed two weeks after each injection, before next immunization if any, to examine sera for anti-spike IgG antibody levels by ELISA. **(B)** Measurement of T cell response against spike protein by quantifying spot forming IFN-γ secreting splenocytes elicited by vaccination. Two weeks after the third injection, splenocytes were collected and stimulated by spike protein. Responsive T cells secreting IFN-γ were counted by IFN-γ ELISPOT assay. (**C, D)** Measurement by ELISA of total IFN-γ **(C)** and IL-2 **(D)** secreted by splenocytes after spike protein stimulation.

Strong immune response is typically associated with the release of pro-inflammatory and pro-immune cytokines such as IFN-γ and IL-2 (46–48). To test this, splenocytes were collected after the final injection and used for spike-specific IFN-γ ELISPOT assay. The cultured splenocytes were also stimulated by the spike protein and the culture media were tested by ELISA for IFN-γ and IL-2 levels. All these indicators of immune response showed higher activities elicited by the miniVec plasmid than the traditional counterpart, while the controls only displayed baseline activities (Figure 8C-D).

### No detectable in vivo immunogenicity or toxicity of miniVec plasmid

The above experiments, in addition to demonstrating enhanced vaccination efficacy of the miniVec system, also addressed the important question of immunogenicity of the miniVec backbone independent of the GOI being delivered. In the control injections of empty plasmids devoid of any GOI, there were no detectable immune response in the animals by the various assays (Figure 8C-D), arguing that the miniVec backbone by itself is not immunogenic in vivo.

We then systematically analyzed the toxicity profile of the miniVec backbone. For acute high-dose toxicity, 400 μg of empty plasmid, either miniVec or traditional, was administered via intramuscular injection to young adult mice divided into separate male and female groups, with uninjected animals serving as control. Body weight changes were measured up to 16 days post injection. No significant difference was observed among the groups (Figure 9A, B). All mice appeared healthy and no overt changes in food consumption, appearance or behavior were observed.

**Figure 9.**
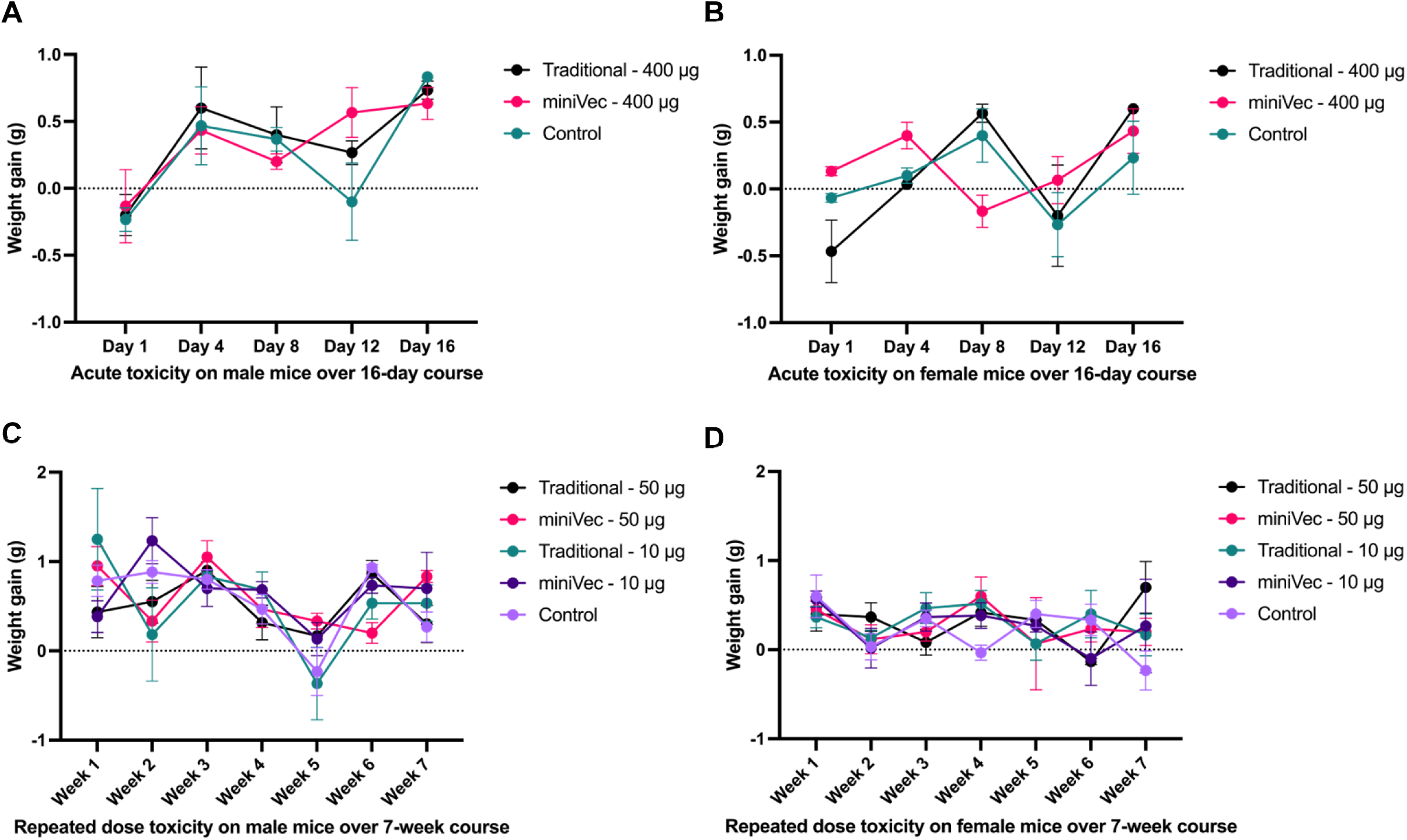
Weight changes of mice administered naked plasmid DNA. **(A, B)** Acute weight changes from a single high-dose administration. One injection of 400 μg of DNA per animal was administered to males **(A)** and females **(B)**, with uninjected mice serving as control (n = 3 each group). Animals were weighed on day 1, 4, 8, 12, and 16 post injection. **(C, D)** Weight changes from repeated-dose administration. A high dose of 50 μg per injection or a low dose of 10 μg per injection was administered twice a week over 4 weeks to males **(C)** and females **(D)**, with uninjected mice serving as control (n = 6 each group). Animals were weighed at the end of each week over 7 weeks after the initial injection.

To evaluate the toxicity of repeated doses, empty plasmids were administered to young adult mice twice a week over 4 weeks, with a high-dose group of 50 μg per injection and a low-dose group of 10 μg per injection. Half of the mice from each group were sacrificed 4 weeks after initial injection to measure organ weight and perform histopathological analysis. The other half were monitored for 7 weeks after initial injection before sacrificed for analysis. Animals were observed daily. No mortality occurred, and animals appeared healthy with no adverse event such as irritation at the injection site, altered food consumption, abnormality in body shape, fur and feces, and changes in behavior. Weight changes were evaluated weekly, which showed no significant difference between the empty plasmids and the uninjected control (Figure 9C, D).

Toxicity was further assessed by hematological measurements, biochemical profile, and organ weight coefficient in both the group sacrificed 4 weeks after initial injection (2 days post final administration) and the group sacrificed 7 weeks after initial injection or 30 days post final administration (Tables S1-S12). Overall, neither the high dose nor the low dose of miniVec plasmid injection displayed any adverse effect, barring some fluctuations that appeared random and dose independent. Meanwhile, histopathological data of liver and kidney did not suggest any significant inflammatory cell infiltrate or necrosis associated with miniVec plasmid administration. One male mouse from the high-dose traditional plasmid group exhibited ballooning degeneration of hepatocytes and vacuolar degeneration in the kidneys (data not shown), suggesting slight toxicity by the traditional plasmid in high dose, though this could not be ruled out as an incidental event.

In sum, naked miniVec plasmid DNA did not trigger any detectable immune response or adverse events from either acute or repeated-dose administration, indicating an excellent safety profile of the miniVec system in vivo.

## DISCUSSION

The miniVec plasmid system represents a significant advancement in vectorology. In particular, it addresses three fundamental requirements for the clinical translation of vectors: manufacturability, functionality especially therapeutic efficacy, and patient safety.

In terms of manufacturability, the miniVec system offers substantial advantages over traditional plasmids. The miniaturized backbone devoid of protein-coding genes lightens metabolic burden on host E. coli, while the small RNA confers constant positive selection to maintain high plasmid copy numbers without antibiotics or any chemical additives. These features result in a massive increase in plasmid yield across different vector designs, and the effect is especially pronounced in large-scale industrial fermentation. Furthermore, the complete elimination of antibiotics and other chemical additives during fermentation simplifies production and QC. These advantages translate into significant cost savings in the manufacturing clinical-grade plasmids by GMP processes. The manufacturing benefits of miniVec also include excellent plasmid stability in E. coli host and excellent plasmid configuration in the form of cccDNA.

In terms of functionality, the advantage of miniVec stems from its minimized prokaryotic sequence that enhances plasmid function in downstream applications. This advantage was consistently demonstrated across multiple experimental systems, including chemical transfection and electroporation, lentivirus packaging, transposon-or CRISPR-mediated genome integration, and in vivo naked DNA transfection and vaccination. Besides increasing clinical efficacy, these improvements translate into compound cost savings stemming from higher yield in viral vector production and reduced dosing, in additional to the cost savings from higher plasmid yield.

The miniVec system also exhibits an excellent safety profile based on both design principles and in vivo studies. The absence of ARGs and a reduction in prokaryotic sequences not only improves manufacturability but also minimizes potential safety risks. In vivo studies in mice confirmed that empty miniVec constructs elicited no immune response, and toxicology analyses revealed no adverse effects on animal health. These features bring miniVec in line with the expectations of best practices from regulatory authorities such as the FDA and EMA.

Other miniaturized plasmid backbone systems have been reported previously (26–29). One notable advantage of miniVec over other systems lies in the fact that, to our knowledge, it is the only plasmid backbone that is completely devoid of any protein coding gene and simultaneously does not require any chemical additive for long-term plasmid maintenance and proliferation in E. coli host.

The miniVec platform presents an especially exciting opportunity for advancing gene and cell therapy. Its demonstrated benefits in diverse applications – from viral vector production, to genome editing tools, to naked DNA vaccination – suggest broad utility across the gene and cell therapy landscape. Importantly, design principles of the system align well with the best practices currently expected by regulatory authorities such as the FDA and EMA for therapeutics manufacturing, particularly regarding the elimination of unnecessary genetic elements.

In conclusion, the miniVec platform represents a major innovation in plasmid DNA technology. Its excellent manufacturability, functionality and safety offer considerable advantages in the development and manufacturing of therapeutics, especially for gene and cell therapy. It also has potential applications in the food industry. We therefore foresee broad adoption of the miniVec system in many biotechnology and biomanufacturing areas.

## METHODS

### Cloning of miniVec plasmids

The plasmids used in this study were first designed and constructed using traditional backbones. The miniVec plasmids were generated based on the corresponding traditional plasmids, with the prokaryotic backbones replaced by the miniVec backbones. The procedure involves amplifying various miniVec backbones and the target insert fragments on different traditional plasmids separately. The whole miniVec plasmids were then assembled using Gibson Assembly. The assembled products were purified using the magnetic AMPure XP beads (Beckman) and electroporated into miniHost competent cells using electroporator (Bio-Rad) with the parameters of 1800 kV, 25 μF, 5 ms. After recovery, cells were plated on antibiotic-free and additive-free LB agar plates and incubated overnight at 37°C. Positive clones were identified through colony PCR and restriction enzyme digestion.

### Plasmids DNA production in laboratory scale

For traditional plasmids, the E. coli DH5α strains transformed successfully with the target plasmids were cultured overnight in 200 mL LB broth supplemented with either 100 g/mL ampicillin or 50 g/mL kanamycin at 37°C. For miniVec plasmids, the miniHost strains containing the target miniVec constructs were cultured overnight in 200 mL LB broth at 37°C without any antibiotics or other chemical additives. The plasmid DNA was extracted using the NucleoBond Xtra Midi EF kit (Macherey-Nagel) according to the manufacturer’s instructions. Each experimental group included three replicates.

### Plasmid DNA production by industrial fermentation

The successfully transformed miniHost E. coli carrying miniVec plasmid or traditional E. coli carrying traditional plasmid were cultured and expanded during the seed train. Once expanded, inoculation into the fermenter with a 2.7-liter working volume was performed. No antibiotics were used in either fermentation to mimic clinical manufacturing settings. Temperature, dissolved oxygen, agitator speed, and pH were controlled in the same manner for both traditional and miniVec plasmids, using parameters from our platform fermentation technology. Harvest time was optimized individually for each plasmid as is typical in industrial fermentation to achieve the best DNA yield. For the overexpression vector we tested, the optimal harvest time for the miniVec and traditional plasmids were 30h and 32h, respectively. Upon harvest, plasmid DNA was extracted and assessed using the same kit as the laboratory production experiment. Each experimental group included two replicates.

### Stability test of miniVec plasmid in miniHost

The miniHost E. coli transformed with miniVec plasmid encoding kanamycin-resistance gene cassette was first cultured in kanamycin containing medium to ensure full resistance in the beginning of the experiment. Each experimental group included two single clones of transformed miniHost E. coli. From Day 1 to Day 15, the miniHost was cultured in antibiotic-free medium and passaged every day. The OD600 was measured before the passage for assessing the doubling speed. The Day 11 and Day 15 samples were diluted and cultured on antibiotic-free agar plate for randomly picking 96 clonal colonies. These colonies were then subjected to culture on kanamycin plate and antibiotic-free plate. Retention rate was calculated based on the colony number on the kanamycin plate over that on the antibiotic-free plate.

### Chemical transfection of plasmids

HEK293T cells were seeded at a density of 2.5×10⁵ cells per well in 12-well plates one day prior to transfection. On the following day, cell confluence was checked, and the wells with 65–75% confluence were used for chemical transfection experiments. Each experimental group included three replicates. Trans-Hi™ transfection reagent (Tribioscience) was used for all transfections according to the manufacturer’s instructions. Dulbecco’s Modified Eagle Medium (DMEM) was pre-warmed in 37°C water bath. Two 1.5 mL microcentrifuge tubes were prepared for each transfection. Both tubes were filled with 50 μL of the pre-warmed DMEM. Tube A was added with 1 μg of plasmid DNA, and Tube B was added with 3 μL of Trans-Hi reagent. The 1 μg of DNA in Tube A contains either 200 ng of traditional vector or equal molar amount of miniVec topped up with empty vector. After mixing thoroughly, the entire volume from Tube B was added to Tube A and pipetted gently. The mixture was incubated at room temperature for 15 minutes. The transfection mixture was then gently added dropwise to the corresponding well of the 12-well plate, followed by incubation at 37°C. At 12–18 hours post-transfection, the culture medium was removed and replaced with fresh, pre-warmed complete medium of DMEM supplemented with 10% fetal bovine serum. At 48 hours post-transfection, cells were imaged using the IX73 inverted fluorescence microscope (OLYMPUS). Cells were then trypsinized, with one-fourth of each sample reseeded into new 12-well plates, and the remaining three-fourths analyzed by flow cytometry using Attune™ NxT Flow Cytometer (Thermo Fisher). At 96 hours post-transfection, cells were again imaged, trypsinized, and analyzed by flow cytometry. The MFI was calculated by multiplying the fluorescence intensity of the fluorescence-positive cells with the positive rate, applicable for all subsequent MFI calculation.

### Electroporation of plasmids

Jurkat cells were cultured and resuspended at a density of 1×10⁶ cells per 1.5 mL centrifuge tube. Either 10 μg of traditional plasmid or equal molar amount of miniVec was added and the total volume was then topped to 300 μL. For the Gene Pulser Xcell electroporation systems (Bio-Rad) settings, the voltage was set at 222 V and the capacitance was 600 μF, using 2 mm electroporation cuvette with exponential protocol. The mixed solution was transferred to the cuvette and was electroporated, followed by resuspension and incubation for another 48 hours. The MFI was calculated as described above.

### Lentiviral vector packaging

The lentiviral vector packaging system and protocol were as previous described (38). Each experimental group included two replicates. The PEG concentration protocol, the titer measurement and flow cytometry analysis of lentiviral vectors were also based on the literature (49). The transduction target cell was modified to HEK293T cell line with an endogenous reference using the BMP2 gene. The primer sequences are listed in Table S13. The instrument for qPCR was QuantStudio 3 (Applied Biosystems) and the master mix with polymerase was TB Green® Premix Ex Taq™ II (Tli RNaseH Plus) (Takara Bio).

### Transpositions using piggyBac (PB) and Sleeping Beauty (SB)

HEK293T cells were seeded into 12-well plates at a density of 2.5×10⁵ cells per well one day prior to transfection. On the following day, cell confluence was checked and the wells with confluence at 65%–75% were used. Trans-Hi™ transfection reagent (Tribioscience) was prepared, and DMEM was pre-warmed in a 37°C water bath. Transfections were performed using a Trans-Hi to plasmid ratio of 3:1 (n = 3). The total amount of plasmid DNA per well was 1 μg in total, including 100 ng of the SB or PB transposon together with 10 ng of SB100X or hyBPase transposase respectively, in miniVec plasmids. Traditional backbone counterparts of equal molar were used for comparison. The remaining amount of DNA was topped with an empty plasmid. The control samples contained the transposon plasmid on traditional backbone alone without the transposase plasmid and also topped with empty plasmid to 1 μg DNA in total. The co-transfection methods of transposon and transposase for both SB system and PB system were based on previously reported protocols (50, 51). Each experimental group included three replicates. At 12–18 hours post-transfection, the culture medium was removed and replaced with fresh, pre-warmed complete medium of DMEM supplemented with 10% FBS. At 48 hours post-transfection, cells were imaged using the IX73 inverted fluorescence microscope (OLYMPUS). Cells were then trypsinized, with one-fourth of each sample reseeded into new 12-well plates, and the remaining three-fourths analyzed by flow cytometry using Attune™ NxT Flow Cytometer (Thermo Fisher). Cell passaging was performed every 2-3 days until a late passage number where the fluorescence disappeared in the control, with cell imaging and flow cytometry analysis prior to each passaging.

### Homology-independent targeted insertion (HITI)

The design of gRNA for HITI on the HEK293T AAVS1 site was as previous reported (52). The design of the HITI donor plasmid was also based on the literature (53). HEK293T cells were seeded into 12-well plates at a density of 2.5×10⁵ cells per well one day prior to transfection, reaching a confluence of 65%–75% on the following day. On the day of transfection, the transfection mixture was prepared with a Trans-Hi to plasmid ratio of 3:1. For each well, a total of 1 μg plasmid DNA was used, consisting of 500 ng of the gRNA/Cas9 vector and 500 ng of the donor vector in the case of traditional plasmid group. The miniVec group was adjusted based on equal molar transfection, topped with empty plasmid. The control group contained the donor plasmid alone without the helper plasmid and also topped with empty plasmid to 1 μg DNA in total. After incubation for 10-15 minutes at room temperature, the transfection mixture was added dropwise to each well. Each experimental group included two replicates. After 12-18 hours, the culture medium was changed. At 48 hours post-transfection, the cells were imaged, and half of the cells were passaged into new 6-well plates for selection with 1 μg/mL of puromycin (InvivoGen), while the remaining half were collected for flow cytometry to assess fluorescence expression. Four days after the puromycin selection, cells were imaged, subsequently cultured and passaged regularly without puromycin. Cells were passaged every 2-3 days, with fluorescence imaging and flow cytometry performed prior to each passage. The passaging was continued for 19 days until the absence of fluorescence in the control group.

### Intravenous (IV) and intramuscular (IM) administration

A total of four 6-8 week old female BALB/c mice (Charles River) were used in each group. Hydrodynamic IV injections were performed as described in the literature (54). In brief, the injection volume was adjusted to 0.1 mL/g body weight, and the injection rate was set at 400 L/s. A total of 10 g miniVec plasmid or equal molar traditional plasmid of luciferase Luc2 plasmid was diluted in D-PBS (Beyotime) based on the mouse body weight and administered via the tail vein. In vivo imaging was conducted on days 1, 2, 3, and 7 post-injection to assess bioluminescence distribution and intensity. IM injections were performed according to previously described protocol (55). Three days prior to plasmid injection, 100 L of 0.25% bupivacaine hydrochloride (Macklin) was injected into the quadriceps muscle. A total of 50 g miniVec plasmid or equal molar traditional plasmid of luciferase Luc2 vector was diluted with D-PBS to a final volume of 70 L and injected into the muscle. In vivo imaging was performed on days 1, 3, 7, 11, and 21 post-injection to evaluate the distribution and intensity of bioluminescence. To measure bioluminescence, the signals were monitored using the Pierce™ D-luciferin potassium salt (Thermo) according to procedures described in the literature (56).

### Toxicology testing

Acute and repeated-dose toxicology was examined based on protocols derived from previous reports on plasmid DNA vaccine and COVID-19 vaccine (57–59). In brief, acute toxicology was studied on three male and three female 6-8 week old BALB/c mice (Charles River) in each group. On day 0, mice in each group received 160 L IM injections of blank, 400 g miniVec or 400 g traditional plasmids which were empty vectors devoid of any GOI. General activity, behavior, morbidity, mortality, and signs of toxicity were monitored daily. Body weight was recorded throughout the 16-day observation period. In the repeated-dose study, six male and six female 6-8 week old BALB/c mice were used in each group. PBS as control, miniVec or traditional plasmids were administered at two dosage levels: a high-dose group (50 g per injection) and a low-dose group (10 g per injection). Each injection was delivered in a volume of 80 L. Injections were performed twice per week for four consecutive weeks, for a total of eight administrations. Half the mice of each group, namely three mice, were sacrificed on day 28 and the remaining were sacrificed on day 56. The mice were monitored daily for general activity, behavior, and clinical symptoms, including signs of morbidity or distress. Body weight was measured once per week. Average body weight per group was calculated to assess trends in weight gain over time. Hematology profile was assessed on day 28 (48 h after the final injection) and day 56 (30 days after the final injection). The following indicators were measured to evaluate cellular composition and hematopoietic function: red blood cell count (RBC), hemoglobin (HGB), hematocrit (HCT), mean corpuscular volume (MCV), mean corpuscular hemoglobin (MCH), mean corpuscular hemoglobin concentration (MCHC), platelet count (PLT), white blood cell count (WBC), neutrophil count, lymphocyte count, monocyte count, eosinophil count, and basophil count. Biochemical profile was also assessed on days 28 and 56 to evaluate liver and kidney functions. The following markers were measured: alanine aminotransferase (ALT), aspartate aminotransferase (AST), alkaline phosphatase (ALP), total bilirubin (T-bil), urea (UREA), creatinine (CREA), total protein (TP), albumin (ALB), glucose (Glu), and total cholesterol (TC). On the day 28 and day 56, the euthanized mice were subject to necropsy. Major organs were examined macroscopically for any pathological changes. The weights of solid organs including the heart, liver, spleen, lungs, kidneys, brain, thymus, muscle, and gonads (testes/ovaries) were recorded, and organ weight coefficient were calculated using the following formula:

Organ weight coefficient (%) = (Organ Weight / Body Weight) × 100%

### Naked DNA vaccination

The study protocol to evaluate naked plasmid DNA as COVID-19 vaccine to trigger immune response against the spike protein was based on previous reports (60, 61). A total of five 9-week old female BALB/c mice (Charles River) were used to inject PBS as control, traditional and miniVec versions of the COVID-19 naked DNA vaccine. Four mice were used to inject empty plasmids of traditional backbone or miniVec backbone. Three days before plasmid DNA injection, mice received an IM injection of 50 L 0.25% bupivacaine hydrochloride solution prepared in saline (Macklin) into the quadriceps muscle. Each mouse then received 100 g plasmid DNA in 80 L PBS, or plain 80 L PBS as control. Immunizations were administered on days 0, 14, and 28. Blood samples obtained via retro-orbital bleeding were collected on days 0, 14, 28, and 42 to examine the immune response, before subsequent administration when applicable. The detection of specific antibody in serum against COVID-19 spike protein was a generic ELISA adapted with recombinant COVID-19 spike protein antigen (Sino Biological) as coated capture agent, and HRP-conjugated goat anti-mouse IgG antibody (Beyotime) as detection antibody for colorimetric measurement. Meanwhile, splenocytes were collected from the euthanized mice on day 42 for IFN-γ ELISPOT assay and cytokines ELISA from the cultured splenocytes supernatant. To measure the COVID-19 spike protein-specific IFN-γ secretion, the mouse IFN-γ ELISPOT assay kit (U-CyTech) was used and assay was performed according to the manufacturer’s instructions. In brief, splenocytes were resuspended and added to the PVDF membrane Multiscreen 96 well plate (Millipore) at 2×10⁵ cells/well, then subjected to stimulation by 2 μg/mL of SARS-CoV-2 Spike Peptide Pools (Sino Biological) while following the typical ELISPOT workflow. Splenocytes were also resuspended and cultured in 24 well plate at 5×10^6^ cells/well while stimulated by 2 μg/mL of the of SARS-CoV-2 Spike Peptide Pools (Sino Biological). The supernatant was collected after 40 hours of culture and subjected to detection of mouse IL-2 and mouse IFN-γ using the corresponding uncoated ELISA kit (Thermo) according to the manufacturer’s instructions.

---

**Table S1.** Hematological profile of male BALB/c mice in repeated dose toxicity test at Day 28. Values are in Mean ± SEM (n=3).

| | WBC ($10^9/L$) | RBC ($10^{12}/L$) | HGB (g/L) | HCT(%) | MCV(fL) | MCH(pg) | MCHC(g/L) | PLT ($10^9/L$) | Neu% | Lym% | Mon% | Eos% | Bas% |
| --- | --- | --- | --- | --- | --- | --- | --- | --- | --- | --- | --- | --- | --- |
| Control - PBS | 4.9 ± 0.3 | 8.5 ± 0.1 | 157.3 ± 5.2 | 41.2 ± 0.5 | 48.3 ± 0.1 | 18.4 ± 0.5 | 381.3 ± 9.0 | 682.0 ± 42.6 | 32.9 ± 1.1 | 63.1 ± 0.4 | 2.7 ± 0.8 | 1.2 ± 0.3 | 0.1 ± 0.0 |
| Traditional - 50 μg | 3.6 ± 0.8 | 7.3 ± 1.5 | 130.7 ± 27.2 | 36.1 ± 7.5 | 49.4 ± 0.1 | 18.0 ± 0.8 | 364.0 ± 16.8 | 668.0 ± 71.0 | 20.1 ± 3.2 | 77.5 ± 4.3 | 1.9 ± 1.0 | 0.4 ± 0.3 | 0.1 ± 0.0 |
| miniVec - 50 μg | 4.8 ± 0.6 | 8.6 ± 0.1 | 160.7 ± 3.7 | 41.7 ± 0.8 | 48.4 ± 0.1 | 18.7 ± 0.2 | 385.0 ± 3.1 | 711.3 ± 25.8 | 24.1 ± 2.1 | 73.3 ± 2.2 | 1.9 ± 0.2 | 0.7 ± 0.1 | 0.0 ± 0.0 |
| Traditional - 10 μg | 5.2 ± 0.6 | 8.6 ± 0.1 | 160.7 ± 2.6 | 41.5 ± 0.8 | 48.4 ± 0.3 | 18.8 ± 0.2 | 387.7 ± 3.2 | 701.7 ± 13.7 | 29.5 ± 1.2 | 67.3 ± 0.7 | 2.4 ± 0.8 | 0.8 ± 0.1 | 0.0 ± 0.0 |
| miniVec - 10 μg | 3.3 ± 0.2 | 8.6 ± 0.1 | 160.0 ± 1.0 | 41.7 ± 0.7 | 48.6 ± 0.1 | 18.7 ± 0.2 | 383.3 ± 3.9 | 698.0 ± 17.6 | 26.5 ± 3.3 | 70.9 ± 2.6 | 1.9 ± 0.7 | 0.7 ± 0.1 | 0.1 ± 0.0 |

**Table S2.** Biochemical profile of male BALB/c mice in repeated dose toxicity test at Day 28. Values are in Mean ± SEM (n=3).

|  | Glu mmol/L | T-bil umol/L | ALT U/L | AST U/L | ALP U/L | TP g/L | ALB g/L | TC mmol/L | CREA umol/L | UREA mmol/L |
| --- | --- | --- | --- | --- | --- | --- | --- | --- | --- | --- |
| Control - PBS | 5.7 ± 0.8 | 1.1 ± 0.3 | 46.0 ± 5.4 | 184.2 ± 12.1 | 113.3 ± 4.7 | 47.3 ± 1.5 | 27.9 ± 1.2 | 2.7 ± 0.1 | 18.5 ± 1.9 | 9.3 ± 0.4 |
| Traditional - 50 μg | 7.0 ± 1.2 | 1.3 ± 0.1 | 55.4 ± 13.9 | 215.0 ± 54.0 | 114.6 ± 2.3 | 44.8 ± 1.1 | 26.0 ± 0.7 | 2.4 ± 0.1 | 26.9 ± 7.3 | 10.5 ± 1.7 |
| miniVec - 50 μg | 8.4 ± 0.7 | 1.0 ± 0.1 | 43.8 ± 9.4 | 175.7 ± 28.7 | 104.1 ± 9.1 | 44.9 ± 0.8 | 26.0 ± 0.7 | 2.6 ± 0.1 | 17.9 ± 0.8 | 11.9 ± 1.5 |
| Traditional - 10 μg | 4.9 ± 0.4 | 1.2 ± 0.1 | 76.3 ± 13.7 | 186.8 ± 33.8 | 114.7 ± 3.2 | 46.1 ± 1.1 | 28.0 ± 0.4 | 2.5 ± 0.1 | 17.6 ± 1.3 | 7.9 ± 0.7 |
| miniVec - 10 μg | 7.9 ± 0.2 | 1.2 ± 0.1 | 44.2 ± 6.9 | 125.3 ± 24.2 | 107.2 ± 3.2 | 44.3 ± 0.8 | 25.5 ± 0.4 | 2.4 ± 0.1 | 16.1 ± 0.8 | 14.3 ± 0.6 |

**Table S3.** Hematological profile of female BALB/c mice in repeated dose toxicity test at Day 28. Values are in Mean ± SEM (n=3).

| | WBC ($10^9/L$) | RBC ($10^{12}/L$) | HGB (g/L) | HCT(%) | MCV(fL) | MCH(pg) | MCHC(g/L) | PLT ($10^9/L$) | Neu% | Lym% | Mon% | Eos% | Bas% |
| --- | --- | --- | --- | --- | --- | --- | --- | --- | --- | --- | --- | --- | --- |
| Control - PBS | 6.6 ± 1.0 | 7.3 ± 0.3 | 138.7 ± 5.2 | 36.2 ± 1.3 | 49.7 ± 0.3 | 19.0 ± 0.4 | 383.7 ± 5.8 | 614.7 ± 14.7 | 14.8 ± 0.8 | 83.0 ± 0.9 | 1.4 ± 0.0 | 0.7 ± 0.2 | 0.1 ± 0.0 |
| Traditional - 50 μg | 5.8 ± 0.1 | 7.9 ± 0.1 | 145.0 ± 6.1 | 38.9 ± 0.8 | 49.6 ± 0.2 | 18.4 ± 0.5 | 371.7 ± 9.2 | 691.7 ± 50.4 | 20.7 ± 2.3 | 75.6 ± 2.7 | 2.1 ± 0.1 | 1.6 ± 0.3 | 0.0 ± 0.0 |
| miniVec - 50 μg | 6.9 ± 0.7 | 7.8 ± 0.1 | 146.0 ± 3.2 | 38.6 ± 0.7 | 49.6 ± 0.7 | 18.8 ± 0.3 | 379.0 ± 1.2 | 659.3 ± 6.8 | 14.4 ± 0.5 | 83.8 ± 0.4 | 0.9 ± 0.1 | 0.7 ± 0.1 | 0.1 ± 0.1 |
| Traditional - 10 μg | 5.0 ± 0.1 | 7.2 ± 0.1 | 132.7 ± 3.8 | 35.4 ± 0.6 | 49.4 ± 0.1 | 18.5 ± 0.3 | 375.3 ± 6.2 | 667.7 ± 23.7 | 19.3 ± 1.6 | 78.6 ± 1.4 | 1.2 ± 0.2 | 0.9 ± 0.2 | 0.0 ± 0.0 |
| miniVec - 10 μg | 4.1 ± 1.2 | 8.0 ± 1.4 | 132.0 ± 10.8 | 39.6 ± 7.3 | 49.4 ± 0.3 | 17.0 ± 1.6 | 345.0 ± 33.8 | 675.3 ± 72.9 | 21.6 ± 1.1 | 76.2 ± 1.5 | 1.3 ± 0.4 | 0.8 ± 0.2 | 0.0 ± 0.0 |

**Table S4.** Biochemical profile of female BALB/c mice in repeated dose toxicity test at Day 28. Values are in Mean ± SEM (n=3).

|  | Glu mmol/L | T-bil umol/L | ALT U/L | AST U/L | ALP U/L | TP g/L | ALB g/L | TC mmol/L | CREA umol/L | UREA mmol/L |
| --- | --- | --- | --- | --- | --- | --- | --- | --- | --- | --- |
| Control - PBS | 8.9 ± 0.7 | 1.7 ± 0.2 | 24.2 ± 2.8 | 99.4 ± 13.8 | 136.4 ± 4.8 | 46.3 ± 0.9 | 29.9 ± 0.3 | 2.3 ± 0.0 | 18.6 ± 1.7 | 9.4 ± 1.0 |
| Traditional - 50 μg | 8.0 ± 0.3 | 1.3 ± 0.2 | 20.7 ± 0.7 | 103.2 ± 12.8 | 147.8 ± 3.1 | 45.9 ± 1.2 | 29.0 ± 0.7 | 2.2 ± 0.1 | 19.6 ± 0.9 | 11.8 ± 0.4 |
| miniVec - 50 μg | 11.6 ± 0.7 | 1.3 ± 0.2 | 24.1 ± 2.4 | 101.1 ± 3.0 | 131.2 ± 1.9 | 44.0 ± 1.1 | 27.7 ± 0.9 | 2.1 ± 0.0 | 18.4 ± 0.9 | 9.8 ± 0.9 |
| Traditional - 10 μg | 9.2 ± 1.1 | 1.8 ± 0.2 | 21.6 ± 1.0 | 111.9 ± 10.5 | 122.6 ± 3.3 | 43.2 ± 2.1 | 26.8 ± 1.2 | 2.0 ± 0.0 | 17.0 ± 1.5 | 10.4 ± 1.0 |
| miniVec - 10 μg | 9.5 ± 0.8 | 1.5 ± 0.2 | 20.2 ± 1.4 | 88.4 ± 9.8 | 133.8 ± 5.8 | 44.2 ± 0.8 | 27.1 ± 0.2 | 2.2 ± 0.1 | 17.2 ± 1.0 | 10.1 ± 0.4 |

**Table S5.** Hematological profile of male BALB/c mice in repeated dose toxicity test at Day 56. Values are in Mean ± SEM (n=3).

| | WBC ($10^9/L$) | RBC ($10^{12}/L$) | HGB (g/L) | HCT(%) | MCV(fL) | MCH(pg) | MCHC(g/L) | PLT ($10^9/L$) | Neu% | Lym% | Mon% | Eos% | Bas% |
| --- | --- | --- | --- | --- | --- | --- | --- | --- | --- | --- | --- | --- | --- |
| Control - PBS | 2.7 ± 0.5 | 8.2 ± 0.1 | 146.7 ± 2.3 | 40.1 ± 0.4 | 48.6 ± 0.2 | 17.8 ± 0.2 | 366.0 ± 4.5 | 692.0 ± 10.0 | 23.6 ± 0.6 | 74.4 ± 0.6 | 1.4 ± 0.1 | 0.6 ± 0.2 | 0.0 ± 0.0 |
| Traditional - 50 μg | 3.3 ± 0.5 | 8.5 ± 0.1 | 153.3 ± 1.7 | 41.1 ± 0.3 | 48.5 ± 0.0 | 18.1 ± 0.1 | 373.3 ± 1.9 | 676.0 ± 20.4 | 24.2 ± 2.5 | 74.6 ± 2.7 | 0.9 ± 0.2 | 0.3 ± 0.1 | 0.0 ± 0.0 |
| miniVec - 50 μg | 2.4 ± 0.4 | 8.9 ± 0.0 | 157.7 ± 1.5 | 43.2 ± 0.1 | 48.3 ± 0.1 | 17.6 ± 0.2 | 365.0 ± 3.8 | 677.7 ± 17.4 | 25.8 ± 4.8 | 72.1 ± 5.0 | 1.2 ± 0.4 | 0.8 ± 0.2 | 0.1 ± 0.0 |
| Traditional - 10 μg | 3.8 ± 0.3 | 8.4 ± 0.2 | 153.7 ± 2.6 | 40.5 ± 1.0 | 48.1 ± 0.0 | 18.2 ± 0.1 | 380.0 ± 2.9 | 674.3 ± 26.5 | 23.2 ± 3.1 | 74.8 ± 3.2 | 1.2 ± 0.2 | 0.8 ± 0.1 | 0.1 ± 0.0 |
| miniVec - 10 μg | 1.6 ± 0.1 | 8.8 ± 0.0 | 155.0 ± 0.6 | 42.8 ± 0.6 | 48.9 ± 0.7 | 17.7 ± 0.0 | 361.3 ± 4.2 | 661.3 ± 16.5 | 26.3 ± 3.7 | 71.7 ± 3.7 | 1.2 ± 0.4 | 0.8 ± 0.1 | 0.1 ± 0.1 |

**Table S6.** Biochemical profile of male BALB/c mice in repeated dose toxicity test at Day 56. Values are in Mean ± SEM (n=3).

|  | Glu mmol/L | T-bil umol/L | ALT U/L | AST U/L | ALP U/L | TP g/L | ALB g/L | TC mmol/L | CREA umol/L | UREA mmol/L |
| --- | --- | --- | --- | --- | --- | --- | --- | --- | --- | --- |
| Control - PBS | 7.8 ± 0.5 | 0.8 ± 0.3 | 53.2 ± 10.3 | 264.7 ± 56.1 | 93.7 ± 2.2 | 39.3 ± 1.3 | 22.9 ± 0.9 | 1.7 ± 0.2 | 12.4 ± 0.9 | 7.3 ± 0.3 |
| Traditional - 50 μg | 5.3 ± 0.0 | 1.0 ± 0.1 | 115.3 ± 4.1 | 332.1 ± 36.4 | 82.0 ± 4.5 | 39.0 ± 1.9 | 23.3 ± 1.7 | 1.8 ± 0.2 | 17.7 ± 4.2 | 7.0 ± 0.1 |
| miniVec - 50 μg | 6.2 ± 0.5 | 1.0 ± 0.2 | 75.3 ± 26.1 | 149.6 ± 44.1 | 81.9 ± 5.0 | 38.6 ± 0.7 | 23.3 ± 0.4 | 1.6 ± 0.1 | 12.6 ± 0.5 | 7.3 ± 0.6 |
| Traditional - 10 μg | 6.2 ± 0.7 | 1.2 ± 0.1 | 39.0 ± 3.3 | 190.0 ± 31.7 | 87.0 ± 5.0 | 40.2 ± 2.2 | 24.4 ± 1.2 | 1.7 ± 0.2 | 15.0 ± 2.7 | 6.6 ± 0.6 |
| miniVec - 10 μg | 7.0 ± 0.6 | 1.0 ± 0.2 | 83.5 ± 4.2 | 170.7 ± 9.0 | 84.8 ± 1.3 | 40.5 ± 0.6 | 23.4 ± 0.1 | 1.8 ± 0.1 | 13.6 ± 1.2 | 8.2 ± 0.5 |

**Table S7.** Hematological profile of female BALB/c mice in repeated dose toxicity test at Day 56. Values are in Mean ± SEM (n=3).

| | WBC ($10^9/L$) | RBC ($10^{12}/L$) | HGB (g/L) | HCT(%) | MCV(fL) | MCH(pg) | MCHC(g/L) | PLT ($10^9/L$) | Neu% | Lym% | Mon% | Eos% | Bas% |
| --- | --- | --- | --- | --- | --- | --- | --- | --- | --- | --- | --- | --- | --- |
| Control - PBS | 4.6 ± 0.2 | 8.6 ± 0.2 | 155.0 ± 4.0 | 41.7 ± 1.3 | 48.6 ± 0.3 | 18.0 ± 0.2 | 370.7 ± 5.5 | 652.0 ± 32.6 | 19.0 ± 1.5 | 79.3 ± 1.4 | 0.8 ± 0.0 | 0.8 ± 0.2 | 0.1 ± 0.0 |
| Traditional - 50 μg | 4.0 ± 0.4 | 8.8 ± 0.1 | 154.7 ± 1.2 | 42.6 ± 0.3 | 48.3 ± 0.1 | 17.5 ± 0.1 | 363.3 ± 0.7 | 637.3 ± 28.1 | 22.0 ± 0.8 | 75.9 ± 0.6 | 1.3 ± 0.4 | 0.7 ± 0.1 | 0.1 ± 0.0 |
| miniVec - 50 μg | 4.6 ± 0.4 | 8.3 ± 0.1 | 150.7 ± 3.0 | 40.2 ± 0.8 | 48.4 ± 0.2 | 18.1 ± 0.1 | 374.3 ± 0.9 | 624.0 ± 13.6 | 23.5 ± 3.1 | 74.7 ± 3.5 | 1.3 ± 0.4 | 0.5 ± 0.2 | 0.0 ± 0.0 |
| Traditional - 10 μg | 4.1 ± 0.2 | 8.6 ± 0.2 | 159.0 ± 1.7 | 41.9 ± 1.0 | 48.8 ± 0.3 | 18.5 ± 0.2 | 379.3 ± 5.4 | 594.7 ± 25.8 | 20.5 ± 3.3 | 78.0 ± 3.3 | 0.9 ± 0.1 | 0.6 ± 0.1 | 0.0 ± 0.0 |
| miniVec - 10 μg | 4.7 ± 0.9 | 8.8 ± 0.2 | 160.3 ± 3.2 | 43.1 ± 0.9 | 48.9 ± 0.2 | 18.2 ± 0.1 | 371.3 ± 1.7 | 585.0 ± 23.6 | 21.1 ± 2.6 | 77.6 ± 2.7 | 0.7 ± 0.0 | 0.5 ± 0.1 | 0.0 ± 0.0 |

**Table S8.** Biochemical profile of female BALB/c mice in repeated dose toxicity test at Day 56. Values are in Mean ± SEM (n=3).

|  | Glu mmol/L | T-bil umol/L | ALT U/L | AST U/L | ALP U/L | TP g/L | ALB g/L | TC mmol/L | CREA umol/L | UREA mmol/L |
| --- | --- | --- | --- | --- | --- | --- | --- | --- | --- | --- |
| Control - PBS | 6.1 ± 0.4 | 1.3 ± 0.1 | 42.8 ± 0.6 | 186.3 ± 8.0 | 80.9 ± 3.5 | 37.5 ± 1.3 | 22.9 ± 0.2 | 1.5 ± 0.1 | 12.4 ± 0.9 | 7.3 ± 0.1 |
| Traditional - 50 μg | 5.8 ± 0.7 | 0.9 ± 0.1 | 73.8 ± 7.5 | 157.0 ± 11.9 | 85.5 ± 11.1 | 35.2 ± 0.9 | 21.9 ± 1.0 | 1.5 ± 0.1 | 13.3 ± 1.0 | 7.2 ± 0.5 |
| miniVec - 50 μg | 4.5 ± 0.4 | 1.0 ± 0.2 | 61.2 ± 2.9 | 175.0 ± 17.8 | 79.0 ± 4.2 | 38.7 ± 1.5 | 23.9 ± 1.3 | 1.5 ± 0.1 | 14.5 ± 1.2 | 7.2 ± 0.6 |
| Traditional - 10 μg | 5.6 ± 0.4 | 1.1 ± 0.2 | 48.7 ± 6.2 | 172.4 ± 36.8 | 78.7 ± 3.8 | 38.1 ± 0.7 | 23.0 ± 0.6 | 1.5 ± 0.1 | 13.4 ± 0.9 | 7.2 ± 0.8 |
| miniVec - 10 μg | 5.8 ± 0.4 | 1.0 ± 0.2 | 70.3 ± 9.5 | 240.2 ± 68.6 | 82.6 ± 5.9 | 39.0 ± 3.0 | 24.2 ± 2.1 | 1.5 ± 0.1 | 19.8 ± 4.9 | 7.7 ± 1.0 |

**Table S9.** Organ weight coefficient of male BALB/c mice in repeated dose toxicity test at Day 28. Values are in Mean ± SEM (n=3).

|  | Heart | Liver | Spleen | Lungs | Kidneys | Brain | Thymus | Muscle injection site | Muscle | Testes |
| --- | --- | --- | --- | --- | --- | --- | --- | --- | --- | --- |
| Control - PBS | 0.56±0.04 | 4.00 ±0.16 | 0.33 ±0.02 | 0.65 ±0.02 | 1.53 ±0.04 | 1.87 ±0.04 | 0.18 ±0.02 | 0.82 ±0.02 | 0.81 ±0.03 | 0.83 ±0.03 |
| Traditional - 50 μg | 0.57 ±0.02 | 4.11 ±0.09 | 0.32 ±0.01 | 0.62 ±0.01 | 1.55 ±0.04 | 1.83 ±0.02 | 0.17 ±0.02 | 0.81 ±0.01 | 0.80 ±0.05 | 0.81 ±0.04 |
| miniVec - 50 μg | 0.49 ±0.01 | 4.31 ±0.15 | 0.34 ±0.01 | 0.58 ±0.02 | 1.67 ±0.10 | 1.82 ±0.02 | 0.15 ±0.00 | 0.81 ±0.03 | 0.84 ±0.00 | 0.80 ±0.04 |
| Traditional - 10 μg | 0.58 ±0.04 | 4.01 ±0.08 | 0.30 ±0.02 | 0.52 ±0.08 | 1.50 ±0.05 | 1.88 ±0.02 | 0.13 ±0.01 | 0.87 ±0.03 | 0.79 ±0.02 | 0.83 ±0.03 |
| miniVec - 10 μg | 0.56 ±0.02 | 4.57 ±0.02 | 0.33 ±0.02 | 0.40 ±0.17 | 1.73 ±0.08 | 1.80 ±0.01 | 0.15 ±0.01 | 0.80 ±0.03 | 0.82 ±0.06 | 0.81 ±0.01 |

**Table S10.** Organ weight coefficient of female BALB/c mice in repeated dose toxicity test at Day 28. Values are in Mean ± SEM (n=3).

|  | Heart | Liver | Spleen | Lungs | Kidneys | Brain | Thymus | Muscle injection site | Muscle | Ovaries and uterus |
| --- | --- | --- | --- | --- | --- | --- | --- | --- | --- | --- |
| Control - PBS | 0.57 ±0.02 | 4.59 ±0.10 | 0.46 ±0.01 | 0.74 ±0.04 | 1.40 ±0.02 | 2.40 ±0.02 | 0.20 ±0.00 | 0.79 ±0.06 | 0.74 ±0.10 | 0.60 ±0.03 |
| Traditional - 50 μg | 0.53 ±0.04 | 4.33 ±0.07 | 0.41 ±0.01 | 0.69 ±0.02 | 1.37 ±0.07 | 2.25 ±0.08 | 0.17 ±0.04 | 0.83 ±0.02 | 0.82 ±0.00 | 0.77 ±0.03 |
| miniVec - 50 μg | 0.54 ±0.03 | 4.04 ±0.23 | 0.44 ±0.03 | 0.74 ±0.04 | 1.19 ±0.01 | 2.22 ±0.05 | 0.24 ±0.03 | 0.80 ±0.02 | 0.77 ±0.01 | 0.59 ±0.08 |
| Traditional - 10 μg | 0.60 ±0.02 | 4.22 ±0.17 | 0.43 ±0.03 | 0.69 ±0.05 | 1.35 ±0.03 | 2.29 ±0.05 | 0.23 ±0.02 | 0.78 ±0.04 | 0.78 ±0.01 | 0.76 ±0.04 |
| miniVec - 10 μg | 0.54 ±0.01 | 4.28 ±0.12 | 0.45 ±0.05 | 0.79 ±0.05 | 1.50 ±0.02 | 2.29 ±0.03 | 0.19 ±0.04 | 0.84 ±0.03 | 0.80 ±0.01 | 0.79 ±0.11 |

**Table S11.** Organ weight coefficient of male BALB/c mice in repeated dose toxicity test at Day 56. Values are in Mean ± SEM (n=3).

|  | Heart | Liver | Spleen | Lungs | Kidneys | Brain | Thymus | Muscle injection site | Muscle | Testes |
| --- | --- | --- | --- | --- | --- | --- | --- | --- | --- | --- |
| Control - PBS | 0.46 ±0.01 | 4.10 ±0.08 | 0.29 ±0.01 | 0.64 ±0.02 | 1.53 ±0.10 | 1.74 ±0.02 | 0.10 ±0.01 | 0.79 ±0.06 | 0.79 ±0.03 | 0.77 ±0.02 |
| Traditional - 50 μg | 0.48 ±0.01 | 4.25 ±0.08 | 0.28 ±0.01 | 0.65 ±0.04 | 1.59 ±0.03 | 1.73 ±0.01 | 0.10 ±0.01 | 0.80 ±0.03 | 0.75 ±0.02 | 0.78 ±0.02 |
| miniVec - 50 μg | 0.49 ±0.02 | 4.10 ±0.14 | 0.30 ±0.00 | 0.67 ±0.03 | 1.59 ±0.08 | 1.76 ±0.02 | 0.11 ±0.01 | 0.82 ±0.06 | 0.84 ±0.02 | 0.84 ±0.01 |
| Traditional - 10 μg | 0.50 ±0.01 | 4.09 ±0.07 | 0.30 ±0.02 | 0.62 ±0.04 | 1.48 ±0.03 | 1.75 ±0.05 | 0.10 ±0.01 | 0.83 ±0.02 | 0.77 ±0.02 | 0.75 ±0.02 |
| miniVec - 10 μg | 0.51 ±0.03 | 4.30 ±0.08 | 0.30 ±0.00 | 0.62 ±0.01 | 1.77 ±0.08 | 1.75 ±0.02 | 0.12 ±0.01 | 0.78 ±0.05 | 0.84 ±0.03 | 0.76 ±0.01 |

**Table S12.** Organ weight coefficient of female BALB/c mice in repeated dose toxicity test at Day 56. Values are in Mean ± SEM (n=3).

|  | Heart | Liver | Spleen | Lungs | Kidneys | Brain | Thymus | Muscle injection site | Muscle | Ovaries and uterus |
| --- | --- | --- | --- | --- | --- | --- | --- | --- | --- | --- |
| Control - PBS | 0.53 ±0.04 | 3.85 ±0.14 | 0.40 ±0.02 | 0.75 ±0.01 | 1.34 ±0.08 | 2.27 ±0.06 | 0.16 ±0.02 | 0.73 ±0.02 | 0.76 ±0.02 | 0.68 ±0.07 |
| Traditional - 50 μg | 0.50 ±0.04 | 3.97 ±0.16 | 0.46 ±0.04 | 0.67 ±0.03 | 1.28 ±0.07 | 2.20 ±0.08 | 0.15 ±0.04 | 0.76 ±0.05 | 0.75 ±0.03 | 0.73 ±0.06 |
| miniVec - 50 μg | 0.45 ±0.01 | 3.92 ±0.07 | 0.44 ±0.02 | 0.68 ±0.07 | 1.30 ±0.04 | 2.21 ±0.04 | 0.16 ±0.02 | 0.78 ±0.06 | 0.75 ±0.06 | 1.06 ±0.54 |
| Traditional - 10 μg | 0.55 ±0.10 | 3.96 ±0.09 | 0.41 ±0.05 | 0.68 ±0.08 | 1.27 ±0.03 | 2.23 ±0.08 | 0.16 ±0.03 | 0.76 ±0.01 | 0.77 ±0.08 | 0.75 ±0.15 |
| miniVec - 10 μg | 0.52 ±0.04 | 4.06 ±0.14 | 0.38 ±0.04 | 0.63 ±0.04 | 1.35 ±0.04 | 2.28 ±0.08 | 0.17 ±0.05 | 0.78 ±0.03 | 0.76 ±0.03 | 0.69 ±0.08 |

**Table S13.**
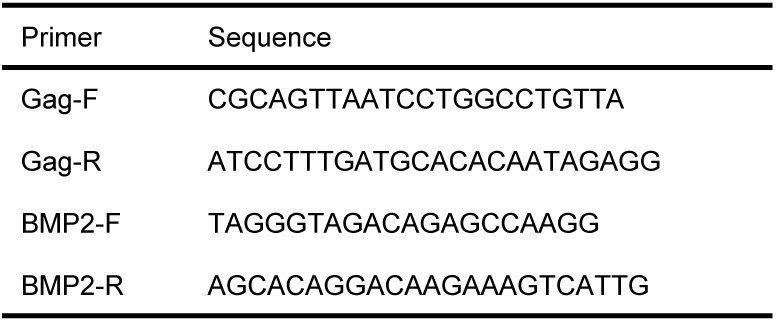
Primers for lentiviral vector titer measurement.

